# TP63, SOX2 and KLF5 Establish Core Regulatory Circuitry and Construct Cancer Specific Epigenome in Esophageal Squamous Cell Carcinoma

**DOI:** 10.1101/825372

**Authors:** Yan-Yi Jiang, Yuan Jiang, Chun-Quan Li, Ying Zhang, Pushkar Dakle, Harvinder Kaur, Jian-Wen Deng, Ruby Yu-Tong Lin, Lin Han, Jian-Jun Xie, Anand Mayakonda, Masaharu Hazawa, Liang Xu, YanYu Li, Luay Aswad, Maya Jeitany, Deepika Kanojia, Xin-Yuan Guan, Melissa J. Fullwood, De-Chen Lin, H. Phillip Koeffler

## Abstract

Transcriptional network is controlled by master transcription factors (TFs) and *cis*-regulatory elements through interacting with target sequences and recruiting epigenetic regulators. By integration of enhancer profiling and chromatin accessibility, we establish super-enhancer (SE) mediated core regulatory circuitry (CRC) for esophageal squamous cell carcinoma (ESCC) and identify tumor cells-dependent CRC TFs-TP63, SOX2 and KLF5. They preferentially co-occupy SE loci and form a positive interconnected auto-regulatory loop through SEs to orchestrate chromatin and transcriptional programming. SE-associated oncogene-*ALDH3A1* is identified as a novel CRC target contributing to ESCC viability. Using circular chromosome conformation capture sequencing (4C-seq) and CRISPR/Cas9 genome editing, the direct interaction between *TP63* promoter and functional enhancers which is mediated by CRC TFs is identified. Deletion of each enhancer decreases expression of CRC TFs and impairs cell viability, phenocopying the knockdown of each CRC TF. Targeting epigenetic regulation by inhibition of either the BET bromodomain or HDAC disrupts the CRC program and its dependent global epigenetic modification, consequently suppressing ESCC tumor growth. Importantly, combination of both compounds result in synergistic anti-tumor effect.

**Graphical Abstract:** 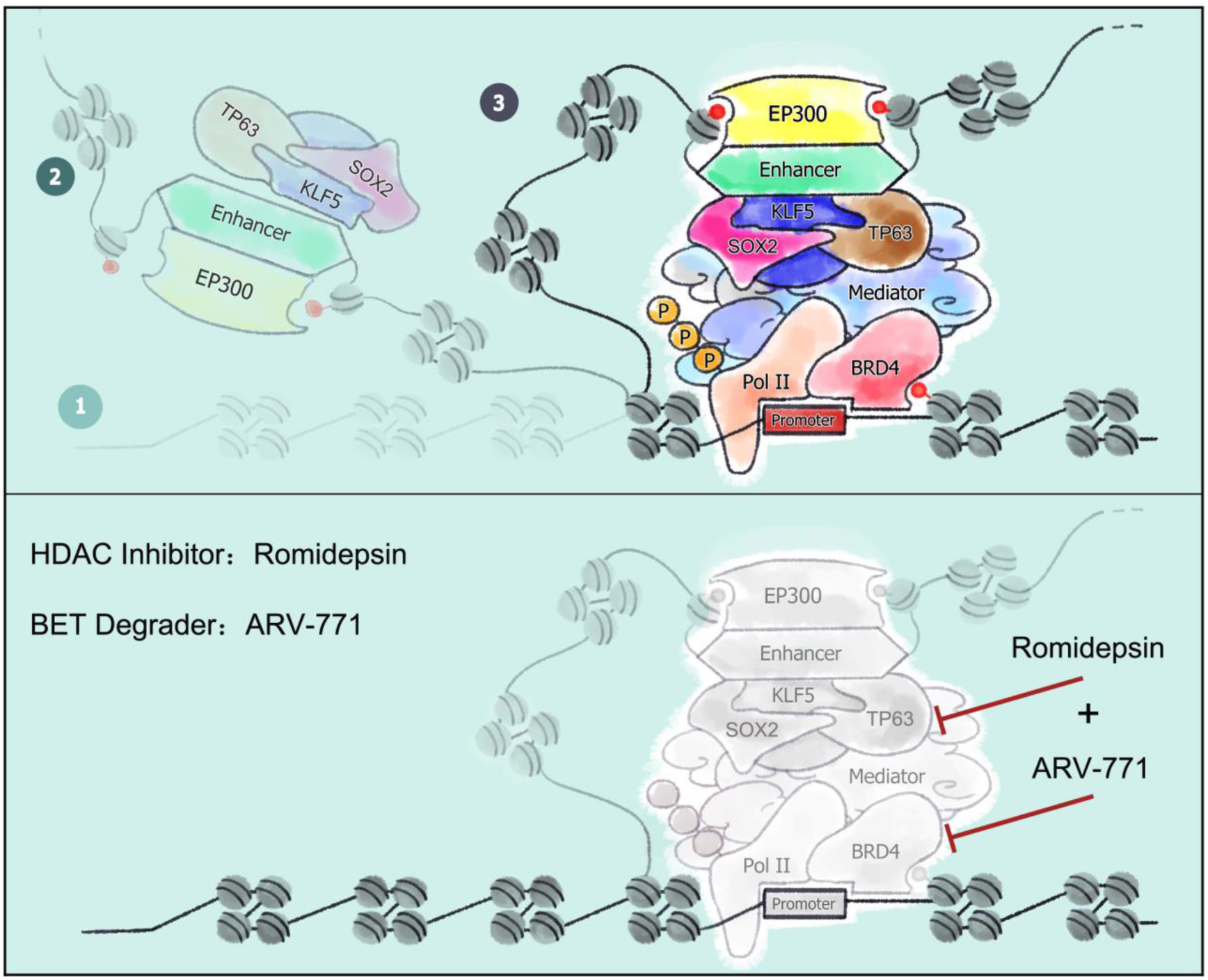

**HIGHLIGHTS:**

- Super-enhancers mediated transcriptional regulatory circuitry is established for ESCC
- TP63, SOX2 and KLF5 as CRC TFs co-localize super-enhancer loci to orchestrate chromatin accessibility and transcriptional dysregulation
- Complex interaction between functional enhancers and *TP63* promoter is mediated by CRC TFs
- *ALDH3A1* is a key downstream target of ESCC CRC and is essential for ESCC cell survival
- BET degrader-ARV-771 and HDAC inhibitor-Romidepsin synergistically inhibit ESCC tumor growth

## INTRODUCTION

Epigenetic characteristics measured by histone modification and chromatin accessibility are useful to delineate the cellular cistrome and transcriptome, which exhibit prominent lineage specificity. These epigenetic characteristics are also notably altered in disease states, such as cancer. In particular, we and others have shown that the clusters of active enhancers called super-enhancers (SEs) mediate transcriptional dysregulation in human cancers (Chipumuro et al., 2014; Hnisz et al., 2013; Jiang et al., 2018; Jiang et al., 2017; Xie et al., 2018; Yuan et al., 2017). These SE loci are occupied by high density of lineage- or cell type-specific transcription factors (TFs), chromatin regulators and coactivators to govern gene expression networks and hence contribute profoundly to cancer biology (Hnisz et al., 2013; Hnisz et al., 2017; Whyte et al., 2013).

In several cell types (e.g., embryonic stem cells, neuroblastoma, medulloblastoma, chronic lymphocytic leukemia and T-cell acute lymphoblastic leukemia) (Boyer et al., 2005; Durbin et al., 2018; Lin et al., 2016; Ott et al., 2018; Sanda et al., 2012), a limited number of TFs (often termed master TFs) bind to their own SEs as well as those of the other members, forming interconnected core regulatory circuitry (CRC) to regulate gene expression of themselves and the other master TFs. Based on these features, CRC TFs can be predicted mathematically by SE mapping coupled with motif enrichment analysis (Saint-Andre et al., 2016). Functionally, CRC TFs orchestrate transcriptional dysregulation in cancer cells by cooperatively enhancing expression of an array of oncogenes via activating their SEs. Notably, this orchestration results in transcriptional addiction in cancer, which can be exploited pharmacologically. However, such a CRC program for esophageal squamous cell carcinoma (ESCC) remains unknown.

ESCC is common (more than 400,000 cases each year) and aggressive malignancy (causing almost 300,000 deaths per year), with a 5-year survival rate less than 20% (Chen et al., 2016; Enzinger and Mayer, 2003). Unfortunately, genome-guided therapeutic strategy is still unavailable for ESCC patients, although ESCC genomic abnormalities have been comprehensively characterized and established (Cancer Genome Atlas Research et al., 2017; Gao et al., 2014; Lin et al., 2018a; Lin et al., 2014; Lin et al., 2018b; Liu et al., 2017; Song et al., 2014). In addition, the high-degree genomic heterogeneity of this cancer further limits application of mutational-targeted therapy. Many findings have suggested that an alternative and appealing therapeutic strategy for a heterogeneous cancer such as ESCC is to target the epigenome (Falkenberg and Johnstone, 2014; Hnisz et al., 2017; Kuhn et al., 2016; Ott et al., 2018). In this regard, a number of epigenetic agents have been developed, including inhibitors against bromodomains and extra-terminal (BET) family proteins and histone deacetylases (HDACs). BET proteins recognize and bind to acetylated lysine residues of histone, and HDACs remove acetyl groups on an array of proteins (Belkina and Denis, 2012; Delmore et al., 2011; Falkenberg and Johnstone, 2014; Filippakopoulos et al., 2010). Both BET and HDAC inhibitors display promising anticancer potential in various types of malignancies through perturbation of multiple components in transcriptional regulation, most notably SEs (Greer et al., 2015; Ott et al., 2018; Ozer et al., 2018; Qu et al., 2017). Specifically, cancer-specific SEs exhibit hypersensitivity to transcriptional inhibition, a primary mechanism underlying the superior anti-tumor potency of BET and HDAC inhibitors (Chapuy et al., 2013; Hnisz et al., 2017; Jiang et al., 2017; Loven et al., 2013; Ott et al., 2018; Yuan et al., 2017).

In this study, we established ESCC-dependent epigenomic-transcriptional regulatory circuitry, determined the exquisite model mediated by core TFs and SEs, and identified crucial downstream targets of CRC program. Destruction of the CRC program by either silencing of CRC TFs, deletion of enhancer elements or administration of epigenetic agents (BET and HDAC inhibitors) potently suppressed ESCC cells proliferation and tumor growth. Strikingly, combinatorial administration of BET and HDAC inhibitors resulted in synergistic anti-ESCC effect.

## RESULTS

### Identification and Characterization of CRC Program in ESCC

Defining epigenomic features is instrumental for understanding gene regulatory programs. Histone markers H3K4me1 and H3K4me3 correlates with active/poised enhancers and active promoters, respectively (Heintzman et al., 2007; Heinz et al., 2015; Rada-Iglesias et al., 2011; Zentner et al., 2011), while H3K27ac (deposited by EP300) separates active from poised regulatory elements (Creyghton et al., 2010; Heintzman et al., 2007; Visel et al., 2009). Using chromatin immunoprecipitation sequencing (ChIP-seq) with antibodies against H3K4me1, H3K4me3 and H3K27ac, the cis-regulatory landscapes of 8 ESCC cell lines was initially defined (Figure 1A). SE-assigned genes were annotated, including well-known SCC oncogenes-*TP63*, *SOX2*, *KLF5*, *CCAT1*, *CTTN, EGFR* (Jiang et al., 2018; Jiang et al., 2017; Watanabe et al., 2014; Xie et al., 2018) (Figure S1A; Tables S1, S2 and S3). We next performed assay of transposase-accessible chromatin with sequencing (ATAC-seq), which showed high concordance with H3K4me1 and H3K27ac signals, as ATAC-seq signals were located at valley regions of H3K4me1 and H3K27ac peaks (Figure 1A), suggesting that transcriptional activity is highly associated with chromatin accessibility.

**Figure 1.**
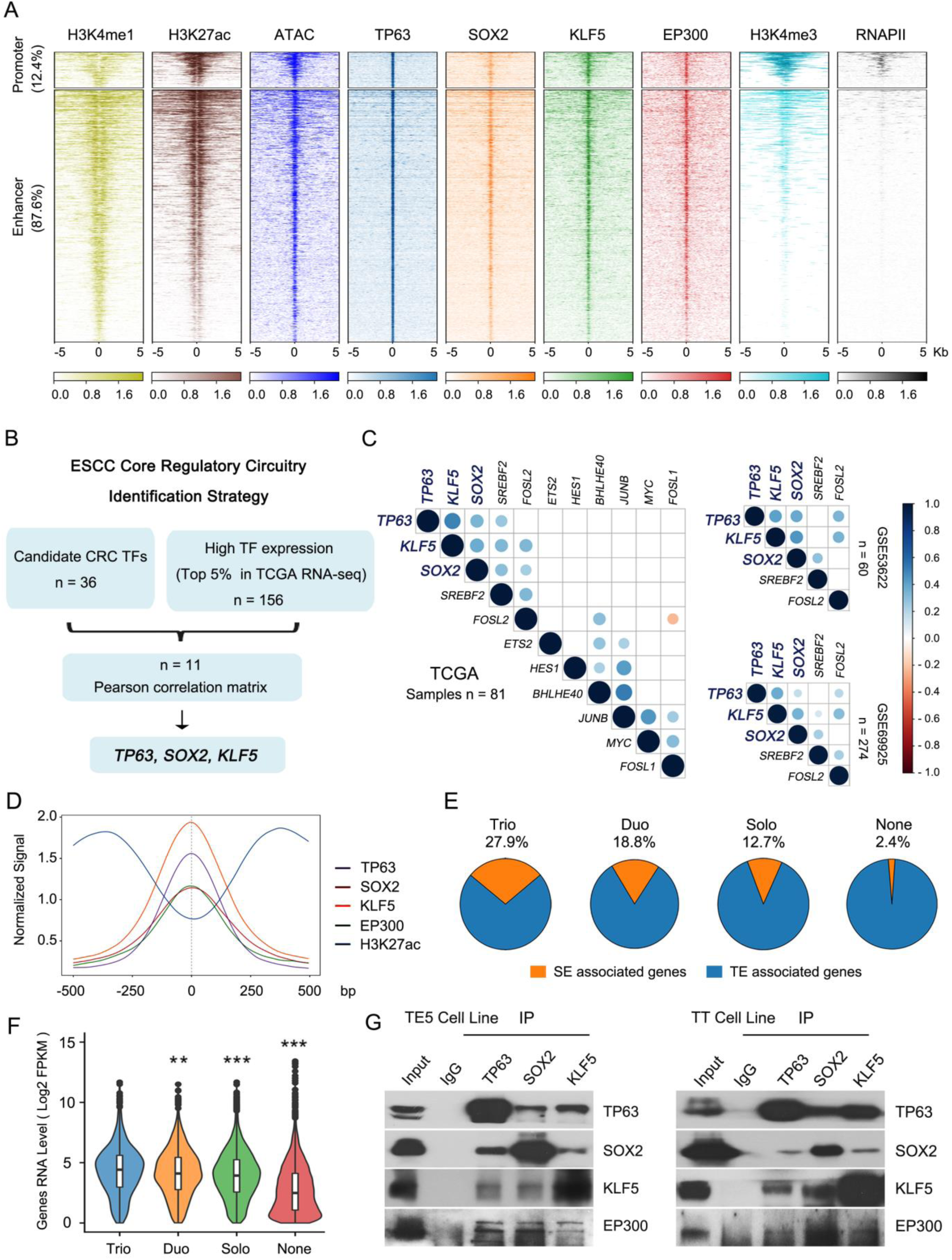
Integrative Analysis Identifies Core Regulatory Circuitry for ESCC. (A) Density plots of the ATAC-seq and ChIP-seq signals of either histone markers or TFs at ± 5 kb windows around the center of TP63 peaks, ranked by TP63 signal in ESCC. Color bars at the bottom show reads-per-million-normalized signals. (B) Schematic diagram of the strategy to identify CRC TFs in ESCC. (C) Pearson correlation matrix for 11 CRC candidates based on TCGA and GEO (GSE53622 and GSE69925) datasets. Color scale is shown on the right. (D) Line plots of indicated ChIP-seq signals centered at the summit of TP63 peaks (± 500 bp). (E) Pie chart illustrating the proportion of SE- and TE-associated genes occupied by trio, duo, solo or none of the three CRC TFs (TP63, SOX2 and KLF5). (F) mRNA expression of genes occupied by trio, duo, solo or none of the three CRC TFs. Error bars, mean ± SD. ** P < 0.01, *** P < 0.001. (G) Co-IP assay of endogenous TP63, SOX2 or KLF5 proteins, followed by western blotting with indicated antibodies to detect the interactions in TE5 and TT ESCC cells.

To construct core regulatory circuitry (CRC) program in ESCC, we first applied CRC_Mapper to these 8 enhancer-annotated ESCC cell lines (Figures 1B and S1A) and identified 36 candidate TFs which occurred in more than 50% at ESCC cell lines (Table S4). Considering that CRC factors are highly expressed in relevant cell types, a total of 156 TFs with high abundance (defined as top 5% of all TFs, TFs were retrieved from TFCheckpoint, http://www.tfcheckpoint.org/) were identified in 81 ESCC samples based on the RNA-seq data from The Cancer Genome Atlas (TCGA) (Table S5). Upon intersection, 11 candidates TFs were identified, including well-established oncogenic factors such as *TP63, SOX2, MYC, SREBF2* (Figures 1B and 1C). Given the co-regulation relationship between CRC members, Pearson correlation analysis of the expression levels of these 11 factors was performed. *TP63*, *SOX2* and *KLF5* consistently had the most significantly positive correlation in 3 different cohorts of ESCC samples (Figure 1C). This result strongly supports recent findings that TP63 directly interacts with SOX2 to co-occupy and regulate hundreds of SEs in SCC cells (Jiang et al., 2018; Watanabe et al., 2014).

As a first step to characterize CRC functions in ESCC, ChIP-seq profiles of TP63, SOX2 and KLF5, along with EP300 and RNAPII were generated and analyzed. Conspicuous trio-binding pattern was observed for the 3 CRC members across the ESCC genome, and the majority of co-occupancy located at active enhancer elements with concordant density of ATAC-seq signals, and surrounded by exceptionally high levels of enhancer markers-H3K4me1 and H3K27ac and coactivator-EP300 (Figures 1A and 1D). Consistent with our previous data (Jiang et al., 2018), the majority of TP63, SOX2 and KLF5 overlapped binding sites located at enhancer regions (87.6%) (Figure 1A). Across the genome, trio-binding (27.9%) was more likely to associate with SEs when compared with either duo- or solo-binding (18.8% and 12.7%, respectively) (Figure 1E). Importantly, SE-associated genes with trio-binding had the highest expression in ESCC cells relative to those with either duo-, solo- or non-binding (Figure 1F), suggesting that the cooperative occupancy among CRC factors produced the highest transcriptional activity.

To determine the mechanistic basis for the cooperation and trio-binding among TP63, SOX2 and KLF5, their protein-protein interaction was examined. Co-immunoprecipitation (Co-IP) revealed that each of TF could pull-down the other 2 proteins, demonstrating the direct interaction among master TFs in ESCC cells (Figure 1G). This result is also consistent with a previous study detecting the direct protein interaction between SOX2 and TP63 in SCCs based on mass spectrometry (Watanabe et al., 2014). In addition, the presence of EP300 protein was observed in either TP63-, SOX2- or KLF5-immunoprecipitated complex (Figure 1G).

Close visualization confirmed prominent trio-occupancy of TP63, SOX2 and KLF5 on their own SE elements as well as others (Figures 2A and S1B), supporting the interconnected auto-regulatory loop involving master TFs and their SEs. Additionally, the ChIP-seq datasets from ESCC cells were compared to those from esophageal adenocarcinoma (EAC) and nonmalignant esophageal mucosa (NEM). Notably, relative to the weaker signals for H3K27ac in EAC and NEM samples, SE elements of TP63, SOX2 and KLF5 were prominently enriched for enhancer markers (H3K4me1 and H3K27ac) and coactivator (EP300) with concomitant accessible chromatin (Figures 2A and S1B), indicating ESCC-specific regulatory network driven by CRC master TFs.

**Figure 2.**
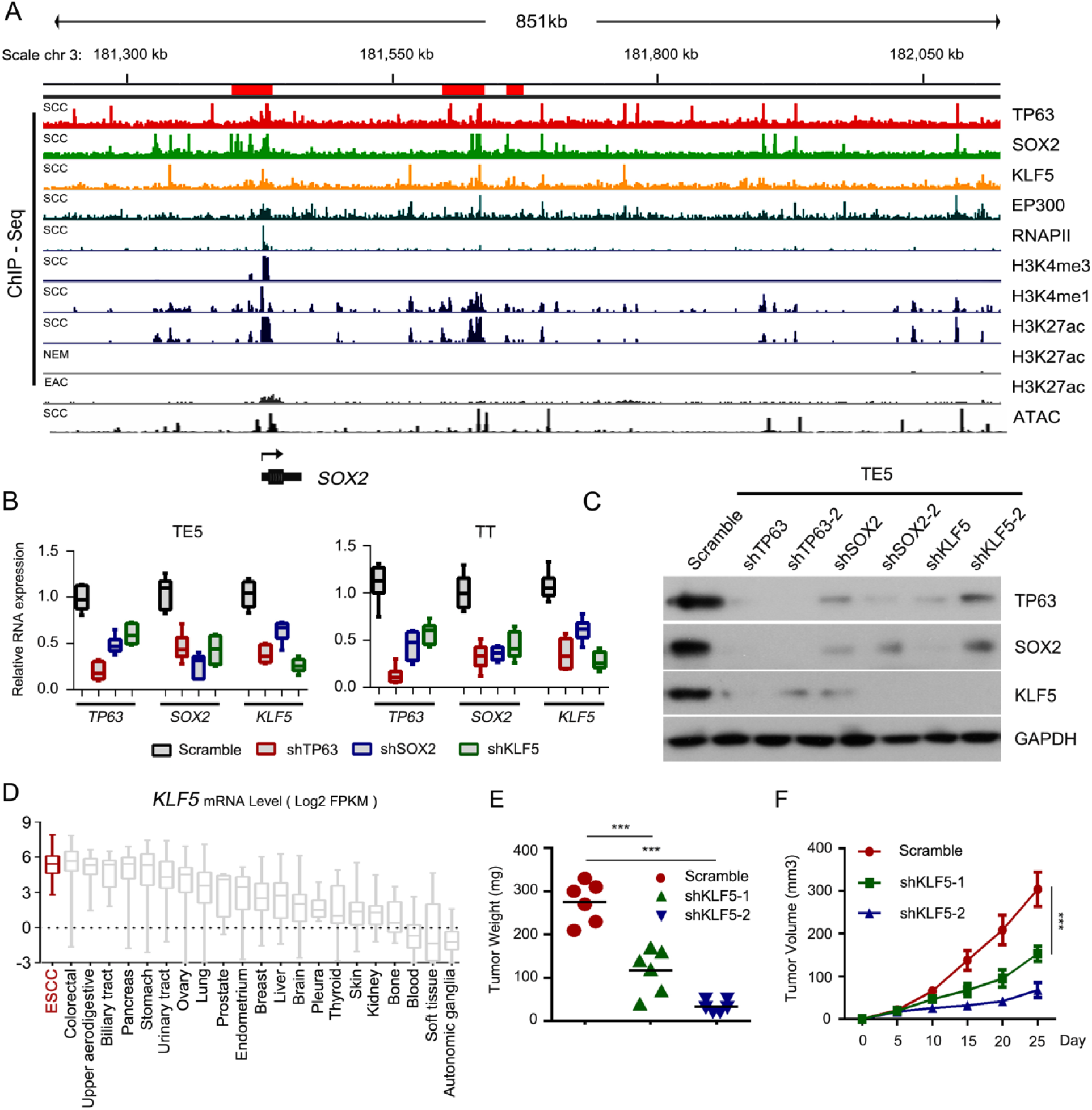
Co-regulatory Circuitry between TP63, SOX2 and KLF5 in ESCC. (A) Integrative genomic viewer (IGV) showing normalized ChIP-seq signals for the indicated antibodies and ATAC-seq tracks at the *SOX2* locus in squamous cell carcinoma (SCC), esophageal adenocarcinoma (EAC) and nonmalignant esophageal mucosa (NEM) samples. Upper red bars denote SE regions. (B) qRT-PCR measuring relative mRNA levels of *TP63*, *SOX2* and *KLF5* upon silencing of each TF relative to scramble controls in TE5 and TT cells. Error bars, mean ± SD. (C) Western blotting analysis showing the protein level of TP63, SOX2 and KLF5 in TE5 cells upon silencing of each of TF using two independent shRNAs. (D) Box plot showing *KLF5* mRNA expression across various types of human cancer cells based on Cancer Cell Line Encyclopedia (CCLE). ESCC cell lines are marked with red. Error bars, mean ± SD. (E) Tumor weights at completion of the study and (F) Growth curves of xenograft volumes in each group. Mice were transplanted with either scramble or silenced KLF5 (two independent shRNAs) TE5 cells. N = 6. Data represent means ±SD, *** P < 0.001.

### Functional Interplay among TP63, SOX2 and KLF5 in ESCC

To dissect the co-regulation of CRC members in ESCC, each of member was silenced using shRNAs, and their expression changes were measured compared with scramble control. Knockdown of any of the TF consistently and significantly downregulated expression of the other two TFs at both mRNA and protein levels (Figures 2B, 2C, S2A and S2B). These data support that ESCC CRC TFs not only co-occupy each other’s SE regions (Figures 2A and S1B), but also form interconnected co-regulatory circuitry.

The functional requirements of TP63 and SOX2 for ESCC have been established (Jiang et al., 2018; Jiang et al., 2017; Xie et al., 2018). However, the biological significance of KLF5 in ESCC is comparably less investigated. The expression of *KLF5* was initially examined, it was top ranked in ESCC samples relative to other types of cancers in either TCGA (Manuscript In Press) or Cancer Cell Line Encyclopedia (CCLE) samples (Figure 2D). This further supports the correlation and co-regulation between KLF5, TP63 and SOX2, since previous results showed that TP63 and SOX2 also had a prominent SCC-specific expression pattern (Jiang et al., 2018; Watanabe et al., 2014). Importantly, downregulation of KLF5 in different ESCC lines markedly suppressed cell proliferation and colony formation (Figures S2C and S2D). KLF5-silenced ESCC cells were transplanted into NSG mice subcutaneously, and consistently, KLF5-depletion diminished tumor growth *in vivo* (Figures 2E and 2F). Western blot and IHC staining of xenograft samples verified that the protein levels of KLF5, TP63 and SOX2 as well as Ki67 (cell proliferation marker) were decreased significantly upon silencing of KLF5 (Figures S2E and S2F). These results confirmed the key pro-ESCC property of KLF5 and further strengthened the inter-regulatory relationship among CRC TFs.

### CRC Factors Cooperatively Promote and Maintain Chromatin Accessibility

To explore whether ESCC CRC members are capable of modulating chromatin accessibility to regulate global gene expression network, ATAC-seq was performed upon silencing each of TF in three different ESCC cell lines (TE5, TT and KYSE140 cells). Strikingly, across the genome, silencing of TP63, SOX2 and KLF5 lost ∼33.9%, ∼47.6% and ∼44.9% of their ATAC-seq peaks, respectively. In contrast, only a very small fraction (∼14.8%, ∼6.5% and ∼9.2% correspondingly) of ATAC-seq peaks gained accessibility (Figures 3A, 3B and S3A-S3D), strongly suggesting that TP63, SOX2 and KLF5 play a crucial role in establishing and maintaining chromatin accessibility in ESCC. This result was also consistent with our earlier finding that TP63 and SOX2 mostly occupy active regulatory regions.

**Figure 3.**
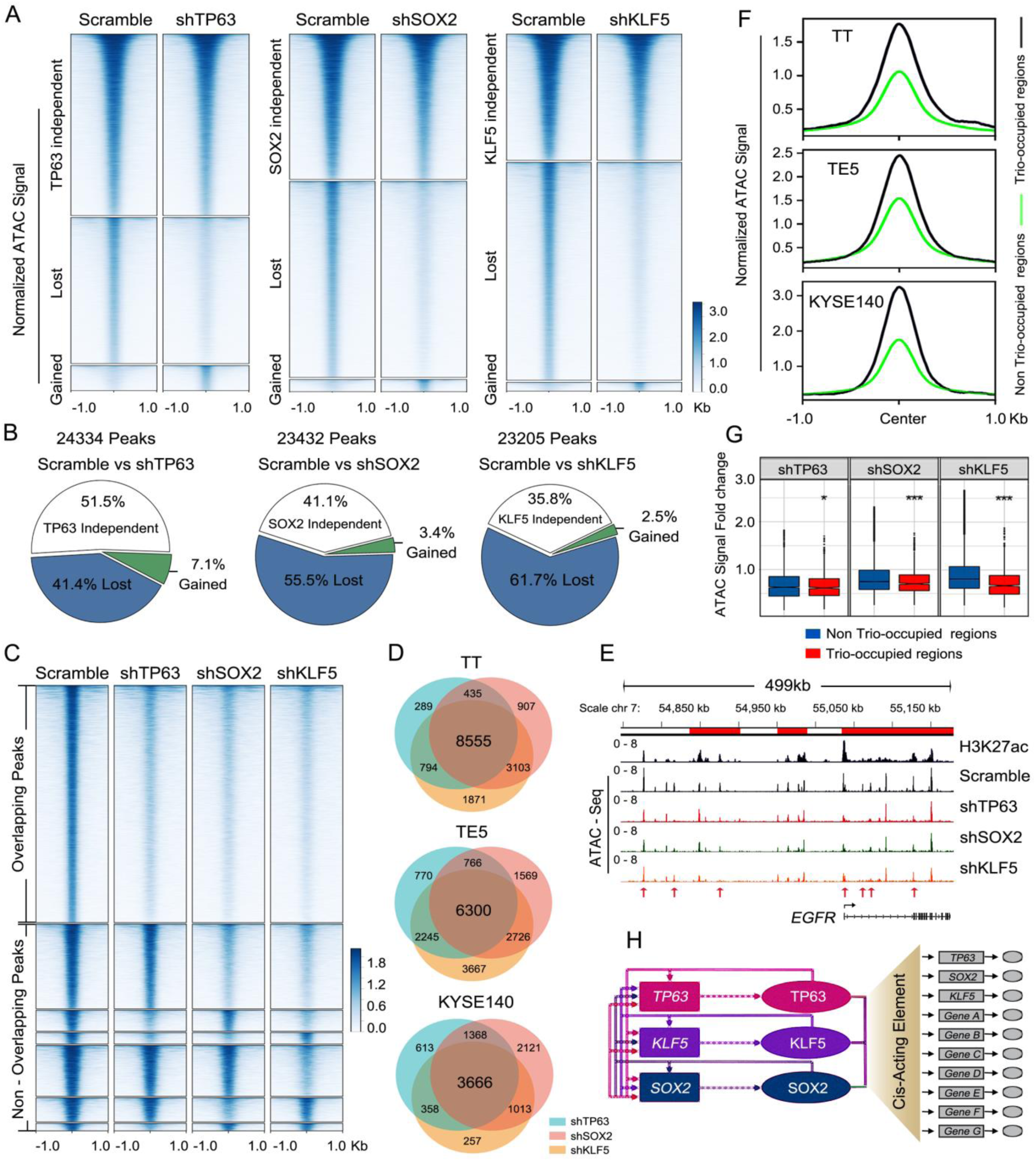
TP63/SOX2/KLF5 Cooperatively Regulate Global Chromatin Accessibility. (A) ATAC-seq heatmaps of differential accessible regions between scramble and CRC TF-silenced TT cells (±1 kb windows flanking the center of summit). Color scale is shown on the right. Heatmaps for KYSE140 and TE5 cell lines are provided in Figures S3A and S3C. (B) Pie charts illustrating proportion of changes in ATAC-seq peaks upon silencing of each of TF in TT cells. These data are derived from panel (A). Pie charts for KYSE140 and TE5 cell lines are provided in Figures S3B and S3D. (C) ATAC-seq heatmaps showing genome-wide lost signals upon silencing of each of TF in TT cells, rank ordered by signal intensity of the scramble group. These regions are further stratified by the degree of overlap among each of the experiment. Lost ATAC-seq signals for KYSE140 and TE5 cell lines are provided in Figures S3E and S3F. (D) Venn diagram of shared and specific genomic regions with lost ATAC-seq signals upon silencing of each of TF across three ESCC cell lines. These data are derived from panel (C) and Figure S3E and S3F. (E) Representative IGV tracks of H3K27ac ChIP-seq and ATAC-seq around *EGFR* locus in either the presence or absence of silencing of TP63, SOX2 or KLF5. (F) Line plots showing higher ATAC-seq signals at TP63/SOX2/KLF5 trio-occupied regions (± 1 kb) than non trio-occupied regions. (G) Box plots of ATAC-seq signals of trio-occupied and non trio-occupied regions upon silencing of either TP63, SOX2 or KLF5. Error bars, mean ± SD, * P < 0.05, *** P < 0.001. (H) Schematic model of CRC program in ESCC.

We next focused on the chromatin regions exhibiting lost of accessibility upon silencing of TP63, SOX2 and KLF5. Strikingly, as much as 34.9% - 53.6% of all lost regions were shared (53.6%, 34.9% and 39.0% in TT, TE5 and KYSE140 cells, respectively) after knockdown of individual TF (Figures 3C, 3D, S3E and S3F), demonstrating that these three factors co-regulate accessibility of thousands of loci along the genome. For example, in an *EGFR* super-enhancer which was recently identified to be co-occupied and transcriptionally co-regulated by TP63 and SOX2 in ESCC (Jiang et al., 2018), multiple enhancer elements were noted with reduced ATAC-seq signals upon silencing of each CRC member (red arrows, Figure 3E). Considering the prominent pattern of trio-occupancy of these master TFs along the genome (Figures 1A and 1D), their ChIP-seq data were integratively analyzed with the dynamic changes in ATAC-seq signals. Not surprisingly, the regulatory elements of trio-occupied by CRC members were overall more accessible (Figure 3F). More importantly, silencing of either TP63, SOX2 or KLF5 led to a more pronounced attenuation of the ATAC-seq signals at trio-occupied regions relative to non-trio-occupied regions (Figure 3G), suggesting that the change of accessibility was a result from their cooperative transcriptional regulation rather than indirect effects. Collectively, these results demonstrate that through direct protein-protein interaction, TP63, SOX2 and KLF5 promote and maintain the chromatin accessibility at thousands of cis-regulatory regions, thereby orchestrating the transcriptional network in a cooperative manner in ESCC cells (Figure 3H).

### *ALDH3A1* Is a Novel Downstream Target of ESCC CRC

We next aimed to identify downstream targets trio-regulated by TP63, SOX2 and KLF5. RNA-seq analysis was performed upon independently silencing of each of TF in ESCC cells (Table S6) (Jiang et al., 2018). Upon integrating RNA-seq results with ATAC-seq data, a total of 175 genes exhibited significant decreased of both mRNA expression (Table S7) and ATAC-seq signals (Figure 4A). These genes included several known TP63/SOX2 targets with prominent functions in SCC, such as *TXRND1* and *CCAT1* (Jiang et al., 2018). Amongst these 175 targets, we were particularly interested in *ALDH3A1* (Aldehyde Dehydrogenase 3 Family Member A1) because it not only showed ESCC-specific expression pattern but also had highest expression in ESCC cells except for *KRT6A* (Figure 4B; Table S7), which encoded Keratin housekeeping protein for keratinization (Hanukoglu and Fuchs, 1983).

**Figure 4.**
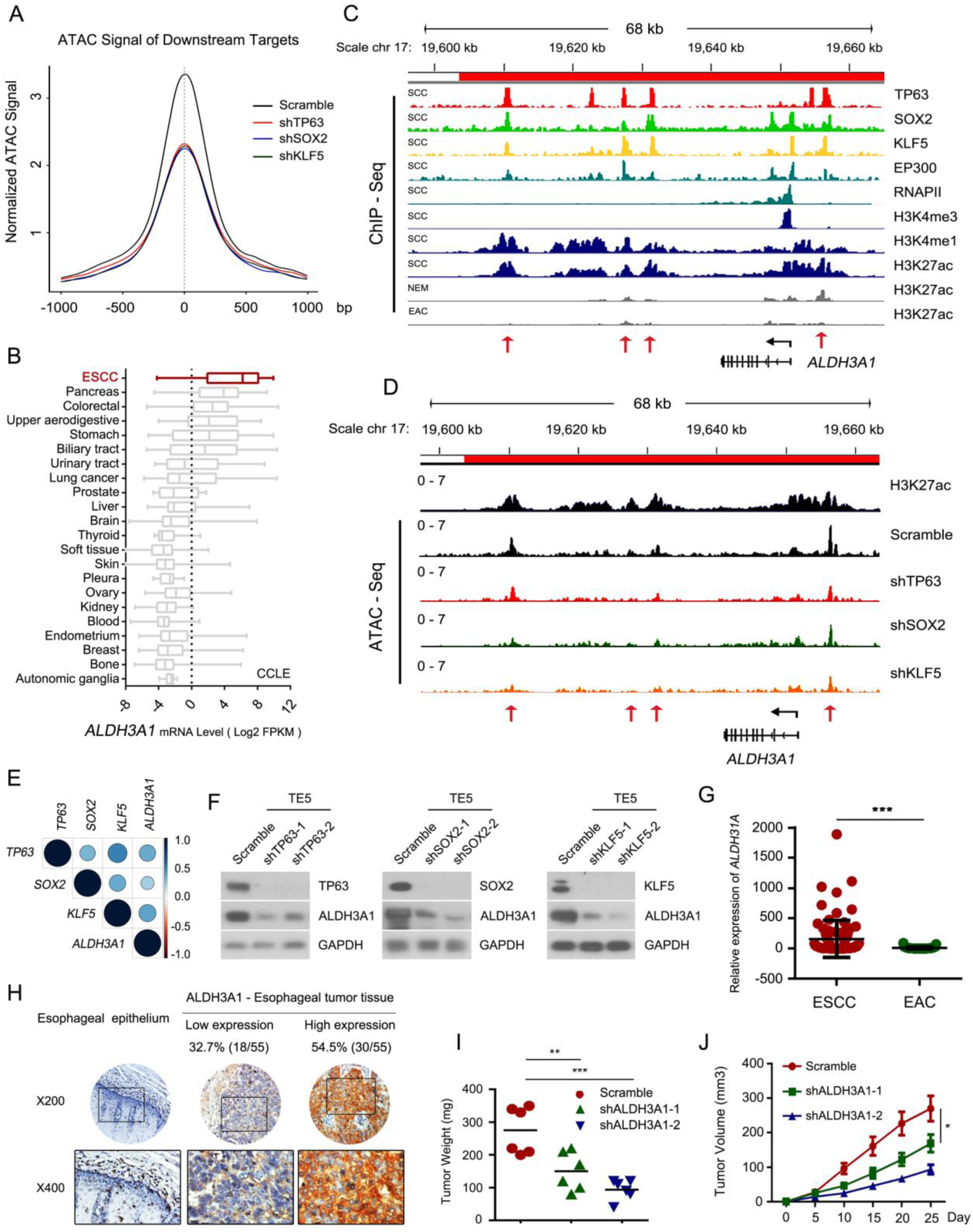
ALDH3A1 is a Key Downstream Target of CRC TFs in ESCC. (A) Line plots showing ATAC-seq signals of downstream targets of CRC TFs. These were decreased upon silencing either TP63, SOX2 or KLF5. (B) Box plots showing mRNA expression of *ALDH3A1* in various cancer cells (CCLE database). ESCC is highlighted in red. (C) IGV tracks showing normalized ChIP-seq of indicated antibodies surrounding *ALDH3A1* gene locus. Red bar denotes SE regions. Red arrows highlight domains trio-occupied by CRC TFs and EP300. (D) ATAC-seq and H3K27ac tracks around the *ALDH3A1* gene locus. Red bar denotes the SE regions. Arrows showing significantly reduced ATAC-seq peaks at the SE occurring after knockdown of either TP63, SOX2 or KLF5. (E) Pearson correlation analysis showing strong positive correlation between CRC TFs and *ALDH3A1* (TCGA RNA-seq data of ESCC samples). (F) Western blotting arrays showing decreased protein levels of ALDH3A1 upon silencing each of CRC TF in TE5 cells. (G) Relative mRNA levels of *ALDH3A1* between ESCC and EAC samples. Data were retrieved from TCGA database. Error bars, mean ± SD, *** P < 0.001. (H) IHC staining of ESCC samples using ALDH3A1 antibody. Upper panel: original magnification, x 200. Bottom panel: zoomed in areas, original magnification, x 400. (I) Tumor weights at completion of the study and (J) Tumor growth curves of scramble and ALDH3A1-silenced groups. N = 6, * P < 0.05, ** P < 0.01, *** P < 0.001.

We interrogated how *ALDH3A1* was regulated by ESCC CRC. ChIP-seq data showed that *ALDH3A1* locus was flanked by broad SE clusters (red bar, Figure 4C), with concomitant open chromatin signals (Figures 4C and 4D). In comparison, the SE of *ALDH3A1* had either a much weaker or undetectable H3K27ac signals in both EAC and NEM samples (Figure 4C), supporting an ESCC-specific regulatory mechanism. The SE constituents of *ALDH3A1* (red arrows in Figure 4C) were prominently trio-bound by all CRC members and EP300 (Figure 4C). More importantly, silencing of any CRC TF led to significant decreased chromatin accessibility at trio-binding regions (Figure 4D). These data suggest a direct regulatory function of CRC TFs on ALDH3A1 transcription, which was further supported by the result that the mRNA level of ALDH3A1 was significantly correlated with each CRC TF (Figure 4E, data from TCGA). Silencing individual CRC members markedly and consistently downregulated expression of ALDH3A1 at both the mRNA and protein levels (Figures 4F, S4A and S4B). Taken together, these results strongly suggest that TP63, SOX2 and KLF5 cooperatively regulate ALDH3A1 transcription, by directly occupying the SE regions of *ALDH3A1* and enhancing their accessibility in ESCC.

Consistent with the enhancer profiles (Figure 4C), expression of *ALDH3A1* was higher in ESCC than EAC samples (Figure 4G). IHC staining of tissue microarray showed strong expression of ALDH3A1 protein in more than half of ESCC tumors (54.5%, 30 of 55) (Figure 4H). In contrast, it was barely detectable in nonmalignant esophagus epithelium. To evaluate the functional significance of *ALDH3A1* in ESCC cells, loss-of-function assays were performed, silencing of ALDH3A1 suppressed cell viability and clonogenic capacity (Figures S4C and S4D). In xenograft assays, knockdown of ALDH3A1 consistently inhibited tumor growth *in vivo* (Figures 4I, 4J and S4E). Taken together, these data identified *ALDH3A1* as a novel and important target of TP63, SOX2 and KLF5, which is specifically upregulated in ESCC and promotes cell proliferation of this cancer.

### Mechanistic Characterization the SE Regions of TP63

To investigate further the mechanism underlying the cooperatively transcriptional regulation among CRC TFs, *TP63* was selected as a study model, which itself was a CRC member and had one of the largest SE regions in several ESCC cell lines (Figures 5A and S1B). Our ChIP-seq data revealed that while *TP63* gene was surrounded by active SEs in ESCC cells (denoted by red bars in Figure 5A), much weaker or almost undetectable H3K27ac signal were present in NEM and EAC (Figure 5A). In addition, most of the SE domains were trio-bound by the three CRC TFs and EP300.

**Figure 5.**
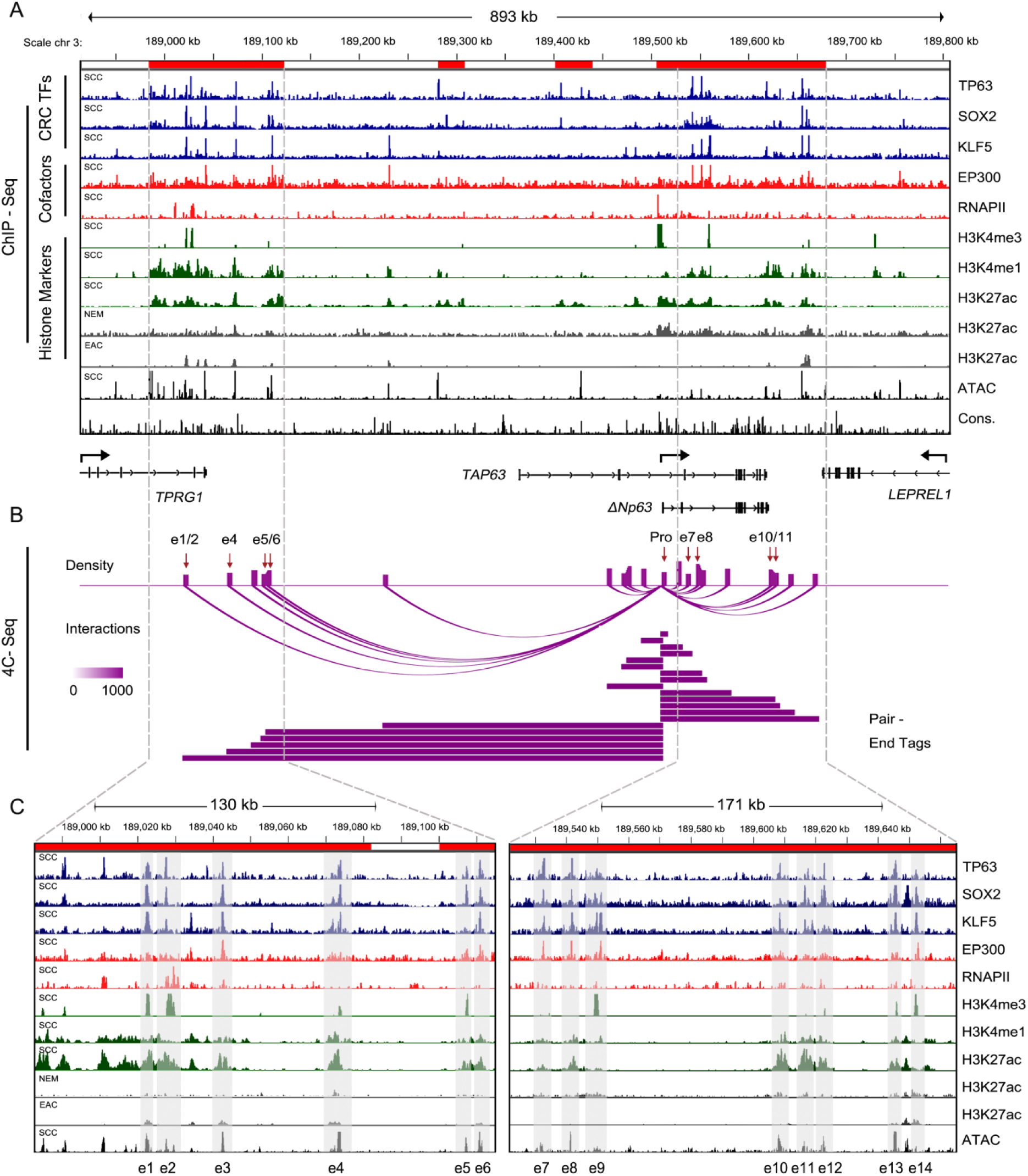
Long-range Interaction between SEs and Promoter of *TP63*. (A) ChIP-seq tracks for indicated antibodies surrounding *TP63* gene locus in ESCC, EAC and NEM samples. (B) Top 20 genomic loci exhibiting long-range interactions with *TP63* promoter in TE5 cells, as identified by circular chromosomal conformation capture (4C)-seq. Vertical and horizontal purple columns indicate interaction density and the distance between interaction loci and *TP63* promoter, respectively. (C) Zoom in view of ChIP-seq tracks as shown in panel (A). Grey shadows highlighting predicted active SEs (separated into e1-e14 enhancer constituents for illustration purpose) which are trio-occupied by CRC TFs and EP300. SE regions are depicted as red bars.

We next employed circular chromosome conformation capture sequencing (4C-seq) assays in TE5 cells to unbiasedly identify DNA regions which had physical contact with the *TP63* promoter (Figure 5B; Table S8). Focusing on the top 20 most significant DNA-DNA interactions (according to *q* value) (Figure 5B), 9 of these 20 regions were located in the enhancer domains (namely, e1, e2, e4-e8, e10 and e11, Figures 5B, 5C and 6A). Considering that only a total of 14 enhancer elements were present within the SE clusters flanking *TP63* promoters (shaded area of bottom panel, Figure 5C), this high-degree of overlap suggests that the majority of this SE was dedicated to activate *TP63* promoter.

A subset of enhancers were also bound by RNAPII and transcribed into non-polyadenylated noncoding RNAs, termed enhancer RNA (eRNA). Importantly, identification of eRNA is a reliable signature of functional enhancers (De Santa et al., 2010; Kim et al., 2010; Li et al., 2016). Hence, expression level of individual enhancers were examined using the *TP63* promoter as a positive control. qRT-PCR detected prominent eRNA expression at e2, e7 and e8 in the ESCC cell lines, which had a comparable or even higher level than present at the promoter region (Figures 6B and S5A). We thus focused on these three SE constituents. Chromosome Conformation Capture (3C) assay followed by Sanger sequencing confirmed chromosomal interactions between these three enhancers and the *TP63* promoter (Interactions were marked by blue shadow of restriction enzyme HindIII in Figures 6C and S5B).

**Figure 6.**
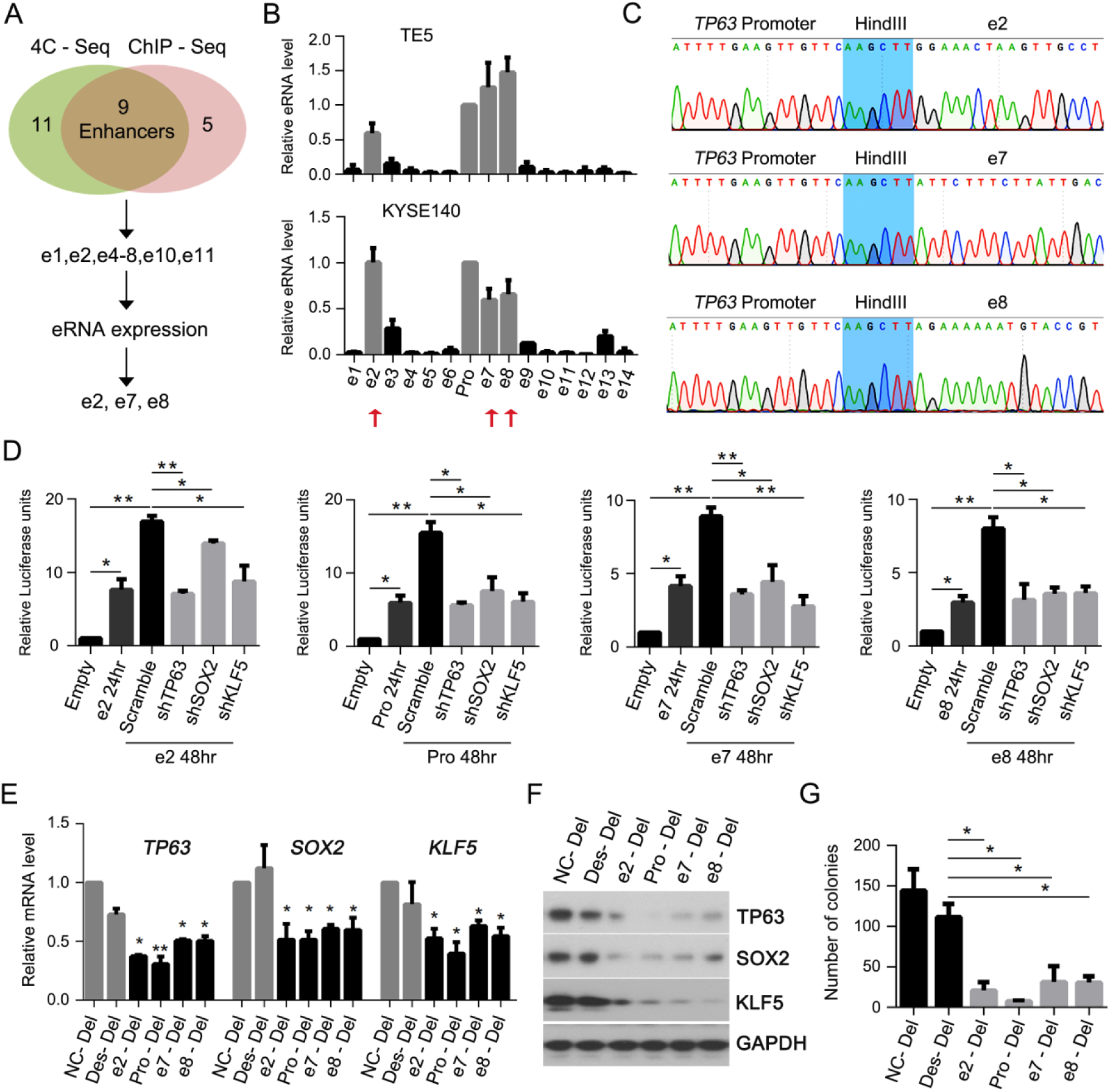
SE Constituents e2, e7 and e8 Contribute to the Activation of *TP63* and Cell Growth of ESCC. (A) Schematic diagram showing the strategy of identifying functional *TP63* enhancers. Upon integrative analysis of ChIP-seq, 4C-seq and enhancer RNA (eRNA) expression (panel B and S5A), e2, e7 and e8 were identified as functional enhancers regulating *TP63* transcription. (B) qRT-PCR measuring eRNA level of individual SE constituents and *TP63* promoter (Pro) in TE5 and KYSE140 cell lines. Similar analysis of the KYSE70, KYSE510 and TT cell lines is provided in Figure S5A. (C) Sanger sequencing results from 3C analysis verifying physical interaction of enhancers (e2, e7 and e8) and *TP63* promoter. Blue shadows are HindIII enzyme sites showing the interaction with *TP63* promoter. (D) Relative activity measured by luciferase reporter assays upon transfection of indicated vectors with including silencing of TP63, SOX2 or KLF5 each separately in TE5 cells. (E-G) Statistical analysis of relative levels of mRNA (E), and protein of CRC TFs (F) as well as colony formation assay (G) upon genomic deletion (Del) of each enhancer element by CRISPR/Cas9. Des (Gene Desert, chr11: 127,292,572-127,294,894) and NC (Negative Control, chr3: 189,368,153-189,370,313) served as negative controls. Data are presented as mean ± SD. Statistical significance (* P < 0.05, ** P < 0.01) were calculated using a two-tailed t test for (D), (E) and (G).

### Disruption of Functional SE Constituents Collapses ESCC CRC Program and Consequently Inhibits ESCC proliferation

To test directly the role of candidate functional SE constituents (e2, e7 and e8) in mediating transcription of *TP63*, luciferase reporter assays were initially performed, and the activity of both the *TP63* promoter and the three enhancers was verified (Figures 6D and S5C). Moreover, the reporter activities of the three enhancers reduced significantly and consistently upon silencing of each CRC member (Figures 6D and S5C), demonstrating that the function of e2, e7 and e8 depends on the activity of CRC TFs.

We next performed CRISPR/Cas9 genome editing and generated cell populations with deletions of either *TP63* promoter or individual SE constituent (validation by Sanger sequencing, Figure S5D). Genomic deletions were also generated in negative control regions, including a gene desert region (Des) and an adjacent region of the *TP63* SE but devoid of H3K27ac signals (NC). Importantly, in comparison with deletion of negative control regions, genomic ablation of either *TP63* promoter or any individual SE constituent caused a significant decrease of expression of both mRNA and protein levels of TP63 (Figures 6E, 6F, S5E and S5F). The expression of SOX2 and KLF5 were concomitantly suppressed (Figures 6E, 6F, S5E and S5F), supporting co-regulation of these CRC factors. Moreover, genomic disruption of either *TP63* promoter or individual SE constituent strongly inhibited proliferation and clonogenic ability of ESCC cells (Figures 6G and S5G), in agreement with the fundamental role of CRC factors in supporting viability of ESCC cells. Notably, in almost all experiments, genomic deletion of any single enhancer element of TP63 produced comparable effects with promoter ablation, highlighting the strong function of these three SE constituents in regulating TP63 expression.

### ARV-771 Evicts BRD4 Protein and Suppresses Proliferation of ESCC Cells

Given the profound role of CRC-mediated transcriptional dysregulation in ESCC cells, the potential anti-ESCC property of BET inhibition was investigated. BET-targeting compounds including competitive BET bromodomain inhibitors and BET-PROTAC degraders (proteolysis-targeting chimera), these have shown promising therapeutic activity in a variety of cancers, particularly in hematopoietic malignancies (Mazur et al., 2015; Ott et al., 2018; Ozer et al., 2018; Qu et al., 2017; Shu et al., 2016; Sun et al., 2018; Zanconato et al., 2018). Initially, expression of BET proteins was examined, BRD2 and BRD4 had much greater abundance than other BET members in TCGA ESCC samples (Figure 7A). ChIP-seq results revealed that *BRD4* was an SE-associated gene, whose SE was enriched with dense H3K27ac, H3K4me1 and ATAC-seq signals (Figure 7B). In contrast, these enhancer peaks were much lower in NEM sample. Moreover, four open chromatin regions were identified (e1-e4 of *BRD4* in Figure 7B) within discrete H3K27ac peaks which were trio-occupied by ESCC CRC members. Importantly, silencing any CRC member substantially decreased expression of BRD4 at both mRNA (Figure S6A) and protein levels (Figures 7C and S6B). We did not observe SE or extensive CRC TF binding in *BRD2* locus (Figure S6C).

**Figure 7.**
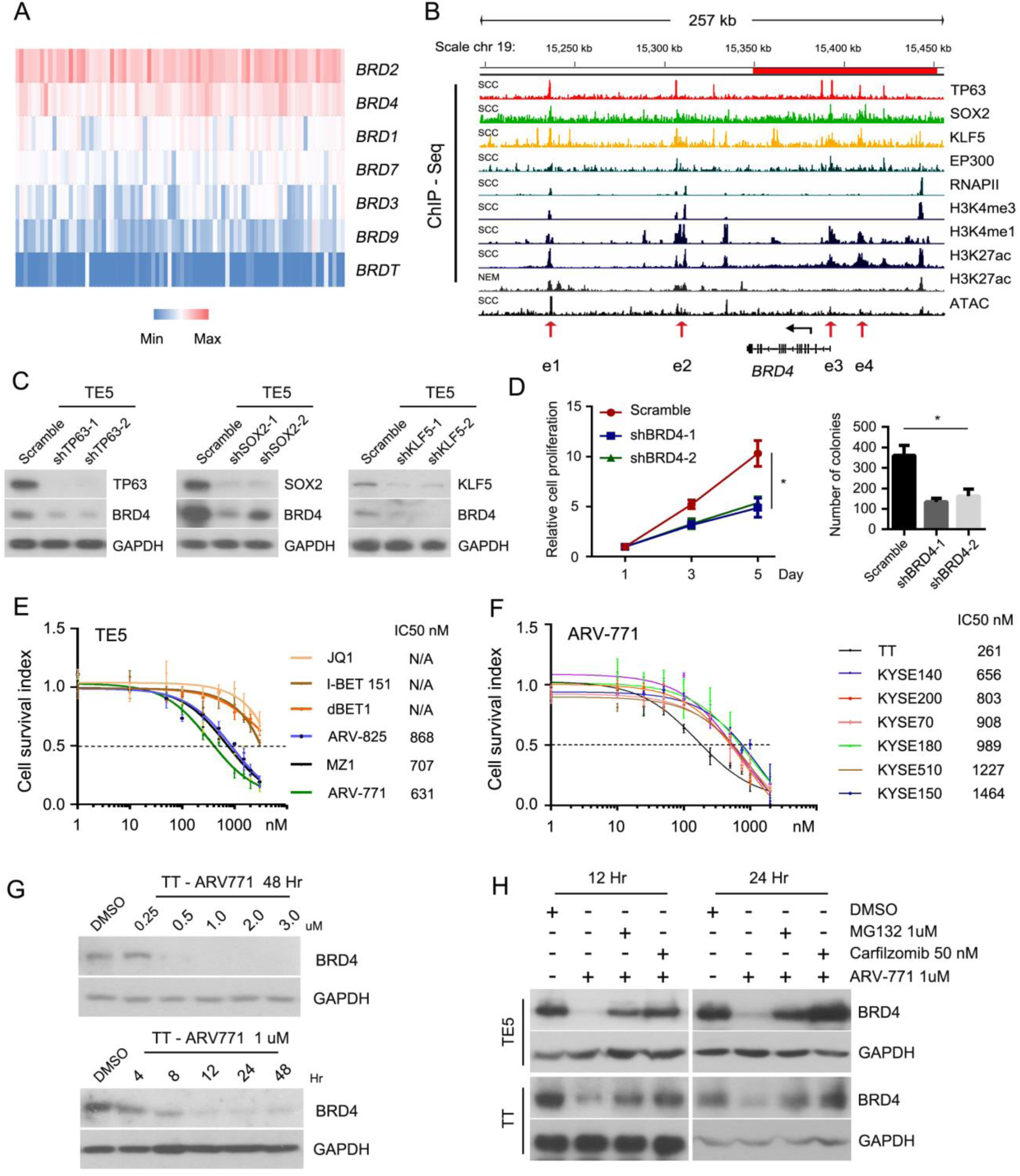
BET Inhibition Potently Suppresses ESCC Cells Growth. (A) Heatmap of expression of BET family members. Data retrieved from TCGA dataset. Deeper red color represents higher expression. (B) ChIP-seq of indicated antibodies and ATAC-seq tracks flanking *BRD4* gene locus. Red bar denotes SE regions. (C) Western blotting analysis for BRD4 and TFs of CRC after silencing of either TP63, SOX2 or KLF5 in TE5 cells. (D) Cell proliferation (Left) and Colony formation (Right) assays following silencing of BRD4 in TE5 cells with two independent shRNAs. Data are mean ± SD from three separate experiments per group. * P < 0.05. (E) Cell growth curves measured by MTT showing sensitivity of TE5 cells to a panel of BET inhibitors and degraders. (F) Dose-dependent curves of all ESCC cell lines during ARV-771 exposure. Data represent mean ± SD of three replicates. (G) Western blotting analysis showing expression of BRD4 in TT cells upon treatment of ARV-771 at different concentrations (Upper) and different durations (Bottom). (H) Western blotting analysis showing protein levels of BRD4 in TE5 and TT cells upon treatment of indicated drugs.

Given the prominent function of BRD4 in other cancer types, we performed a series of loss-of-function assays to explore BRD4-mediated cellular effects in ESCC. Cell proliferation and colony formation ability were significantly suppressed upon silencing of BRD4 (Figures 7D and S6D-S6F), confirming that BRD4 is essential for ESCC growth. Several small-molecule compounds targeting BET proteins in ESCC cells were tested. MTT assay showed that the BET-PROTAC degraders (including ARV-771, MZ1 and ARV-825) were the most potent compounds (Figure 7E). Among of them, ARV-771 exhibited submicromolar IC50 in most ESCC cell lines (Figures 7E and 7F). As expected, ARV-771 inhibited expression of BRD4 both dose- and time-dependently (Figure 7G). Because ARV-771 degrades BET proteins through ubiquitin-proteasome system, this BET-targeting effect was abolished by either MG132 or Carfilzomib proteasome inhibitors (Figure 7H).

### Synergistic Effect between BET Degrader and HDAC Inhibitor in ESCC

In addition to targeting epigenetic readers such as the BET family, inhibition of epigenetic eraser-histone deacetylases (HDACs) can have potent anti-neoplastic activity (Bolden et al., 2006; Ellis et al., 2009; Falkenberg and Johnstone, 2014). Recently, with the advanced understanding of noncoding genome, the effect of HDAC inhibitors on enhancer function was elucidated (Mishra et al., 2017; Sanchez et al., 2018). Intriguingly, pharmacological screening found that HDAC inhibitors reduced TP63 stability in an ubiquitin proteasome-dependent manner (Napoli et al., 2016). Considering that TP63 and its associated CRC program have a profound role in orchestrating ESCC transcriptome, six HDAC inhibitors were tested against ESCC cells. Three of them showed exceptional potency (IC50s: 22-88 nM), with Romidepsin displaying the most potent activity against all of the ESCC cell lines (Figures 8A and S7A). Western blot showed that Romidepsin amongst all HDAC inhibitors was most powerful in degrading TP63 (Figure 8B). Indeed, Romidepsin at as low as 10 nM completely abolished TP63 expression (Figure 8C). Importantly, protein levels of the other two CRC TFs (SOX2 and KLF5) were concomitantly diminished (Figure 8C), again validating co-regulation and co-dependency of these CRC members. Addition of either MG132 or Carfilzomib recovered the protein levels of TP63, as well as those of SOX2 and KLF5 (Figures 8D and S7B), confirming that Romidepsin degrades TP63 protein through an ubiquitin-proteasome system.

**Figure 8.**
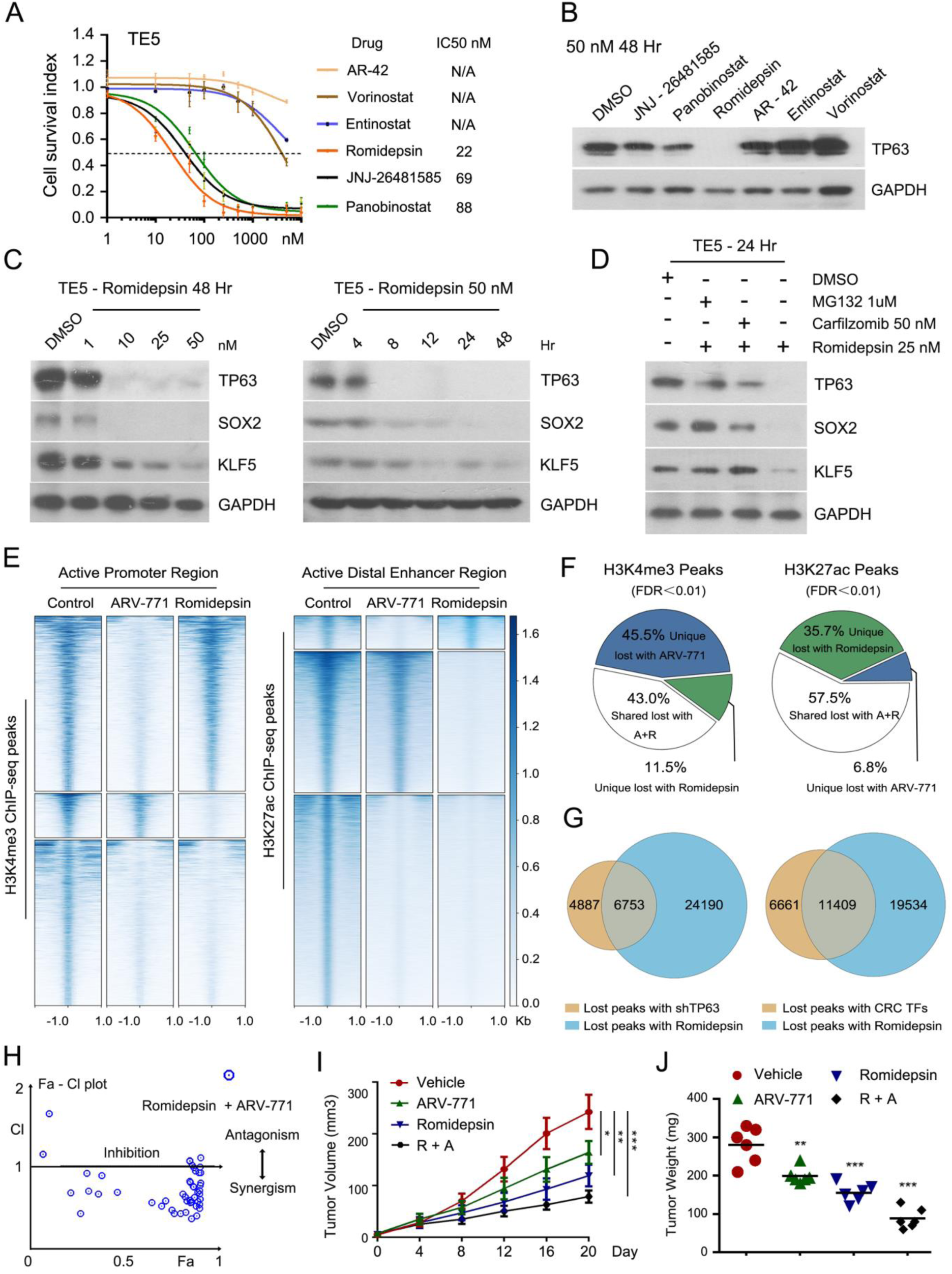
HDAC Inhibition Disrupts CRC Program and Causes a Synergistic Suppression of ESCC Growth when combined with BET Degrader. (A) Cell growth curves testing a panel of HDAC inhibitors. Data are presented as mean ± SD of three replicates. (B) Western blotting showing the protein level of TP63 in TE5 cells upon exposure to various HDAC inhibitors (50nM for 48 hr). (C) Western blotting analysis of expression of CRC TFs treated with either Romidepsin or DMSO at indicated concentrations (Left panel) and time points (Right panel). (D) Western blotting analysis showing protein levels of CRC TFs in TE5 cells upon treatment of indicated chemicals. (E) Heatmaps showing changes of ChIP-seq signals upon treatment of either ARV-771 or Romidepsin in TT cells. Color scale is shown on the right. (F) Pie chart illustrating altered proportion of H3K4me3 and H3K27ac peaks upon treatment of either ARV-771 or Romidepsin in TT cells. Data were derived from panel (E). (G) Lost ATAC-seq signals induced by silencing of either TP63 or a CRC member were shared with lost of H3K27ac signals upon administration of Romidepsin. (H) Fa-CI plots showing synergistic inhibition of ARV-771 and Romidepsin using TT cells, calculated according to results of Figure S7E. CI < 1, = 1, and > 1 indicate eitherdrug synergistic, additive, and antagonistic effect, respectively. (I and J) Tumor growth curves (I) and Tumor weights (J) of mice treated with either vehicle control, ARV-771, Romidepsin or combination treatment of both drugs. Data represent means ± SD, N = 6. * P < 0.05, ** P < 0.01, *** P < 0.001.

Because of the potent anti-neoplastic effect of ARV-771 and Romidepsin in ESCC cells, we sought to understand their mechanisms of actions. Speculating that these inhibitors would disrupt the global epigenetic regulation, H3K27ac and H3K4me3 ChIP-seq were performed in either the presence or absence of these two chemicals. Indeed, treatment of either inhibitor altered the epigenetic modifications across thousands of genomic loci [29,952 (59.6%) H3K27ac and 6,069 (22.5%) H3K4me3 peaks for ARV-771; 48,932 (81.3%) H3K27ac and 19,247 (46.1%) H3K4me3 peaks for Romidepsin, Figure S7C]. Notably, focusing on lost peaks, ARV-771 had a more profound impact on H3K4me3+ promoter regions while Romidepsin showed greater effect on H3K27ac+ distal enhancers (Figures 8E and 8F). On the other hand, 43.0% (H3K4me3) and 57.5% (H3K27ac) of all the “lost regions” were shared between treatments by these two chemicals. These analyses together suggest that ARV-771 and Romidepsin extensively suppress the activity of cis-regulatory elements in ESCC cells. Importantly, these drugs not only co-targeted the epigenetic modification at thousands of shared noncoding regions (white fraction in Figure 8F), but also had specific and complementary effects on a multitude of enhancers and promoters (blue and green fractions in Figure 8F). Moreover, by integration of ATAC-seq data, we observed that 6,753 (58.0%) regions with lost accessibility elicited by silencing of TP63 also showed lose of H3K27ac signals upon administration of Romidepsin (Figure 8G). Consistently, 11,409 (63.1%) loci with lost ATAC-seq peaks upon the depletion of CRC members overlapped regions with lost intensity of H3K27ac following Romidepsin treatment. This strong overlap highlights that Romidepsin suppresses the activity of cis-regulatory elements through reducing expression of TP63 and CRC members.

Considering that ARV-771 and Romidepsin had both shared and unique effects on the epigenomic regulation of ESCC, we hypothesized that a synergistic effect might exist between these two chemicals. Notably, combinatorial treatment with ARV-771 and Romidepsin synergistically inhibited ESCC cell viability (Figures 8H, S7D and S7E) and substantially reduced their IC50s (from 631 nM to 50 nM for ARV-771, from 22 nM to 5 nM for Romidepsin) (Figures 7E, 8A and S7E).

Their synergistic activity was further tested *in vivo* using an ESCC-derived xenograft model. Tumor volume and weight were significantly reduced with either ARV-771 (30 mg/kg) or Romidepsin (1.5 mg/kg) alone compared with vehicle control (Figure 8I and J). Mice were treated with the combination of the two compounds but at half of their original doses (ARV-771 at 15 mg/kg and Romidepsin at 0.75 mg/kg). Importantly, further reductions in tumor growth and tumor weight were observed when compared with single agent treatment (Figures 8I and 8J). IHC staining of the xenografts demonstrated decreased expression of Ki67, BRD4 as well as all three CRC TFs (Figure S7F). Taken together, combinatorial inhibition of BET and HDAC potently and synergistically inhibited ESCC tumor growth both *in vitro* and *in vivo*, through suppressing the activity of cis-regulatory elements as well as CRC-mediated transcription.

## DISCUSSION

By mapping the cis-regulatory landscapes and integrating expression profiles, we established ESCC-specific core regulatory circuitry (CRC) program (TP63, SOX2 and KLF5) that contributes to the ESCC tumorigenesis. ChIP-seq datasets showed that these TFs trio-occupied at the same enhancers and SEs across the ESCC genome, which were marked with dense H3K27ac, H3K4me1, EP300 and ATAC-seq signals. Consistently, enriched binding of KLF5 has been observed in EP300- and H3K27ac-positive regions in head and neck squamous cell carcinoma (HNSCC) (Zhang et al., 2018), indicating a similar epigenetic pattern shared by different types of squamous cell carcinomas (SCCs).

Consistent with the CRC model, TP63, SOX2 and KLF5 trio-bound to the SE elements of their own and each other’s, forming an interconnected transcriptional network in an ESCC-specific manner. Importantly, silencing of any single TF collapsed the entire CRC program, which further led to reduced accessibility at thousands of chromatin elements in ESCC cells. SOX2-containing CRC (together with NANOG and OCT4) has been well-characterized in embryonic stem cells (Boyer et al., 2005; Whyte et al., 2013). In gastric cancer, KLF5, GATA4 and GATA6 are co-regulated by physical interaction on their respective promoters (Chia et al., 2015). These findings suggest that the same TFs can be incorporated into different CRC programs in different cell types to regulate cell-type-specific gene expression patterns. A key characteristic of CRC model is the formation of a feed-forward loop connecting master TFs with the SEs of each CRC member. Here, we investigated in depth how this loop controls SE-promoter interaction at the *TP63* locus. 4C-seq and 3C assays together with CRISPR/Cas9 mediated genome editing showed that three SE constituents (e2, e7 and e8) interacted directly with the *TP63* promoter and contributed to the transcription of TP63, as well as the other master TFs (SOX2 and KLF5). These three enhancer elements were also controlled by CRC TFs, forming a feed forward transcriptional loop. Functionally, deletion of any single SE component potently inhibited ESCC cell growth. Notably, the SE component e2 was recently shown to be engaged in a long-range interaction with the *TP63* promoter through a 3C array in squamous-like pancreatic cancer cells (Andricovich et al., 2018), strongly supporting our unbiased 4C-seq results. Moreover, the other two functional elements (e7 and e8) were also identified previously to be occupied by TP63 itself. Specifically, TP63 binding stimulated the activity of e7 and e8, contributing functionally to the transcription of *TP63* (Antonini et al., 2006; Antonini et al., 2015), congruent with our results (Figures 6 and S5). These findings together highlight the functions of these three SE constituents in the transcriptional activation of TP63 and consequently the entire CRC activity in ESCC.

Through integrative analysis of RNA-seq, ChIP-seq and ATAC-seq, we identified a novel and key direct target of ESCC CRC program, *ALDH3A1*. Notably, *ALDH3A1* also had a strong cluster of ESCC-specific SEs, which were absent in either EAC or NEM samples. Knockdown of any master TF caused a dramatic reduction in ATAC-seq signals at trio-occupied SE regions of *ALDH3A1* by CRC members, confirming their essential role in the direct upregulation of ALDH3A1. Functionally, ALDH3A1 was required for viability of the ESCC cells both i*n vitro* and in *vivo*. As an isoenzyme of aldehyde dehydrogenase superfamily, *ALDH3A1* contributes to tumorigenesis and chemoresistance of cancer cells through detoxifying various aldehydes from both endogenous and exogenous sources by oxidizing aldehydes to the corresponding acids (Kim et al., 2017; Parajuli et al., 2014; Wu et al., 2016). Although the functional role of ALDH3A1 in ESCC was unknown hitherto, large-scale GWAS studies have implied an association between the aldehyde-oxidizing pathway and ESCC susceptibility. Specifically, polymorphic variants of *ALDH2* (associated with changes from Glu487 to Lys487) were found to induce flushing, and they were enriched in individuals of East Asian origin. Notably, these variants increased the risk for ESCC development particularly when these carriers consumed alcohol (Abnet et al., 2018; Brooks et al., 2009; Yokoyama et al., 2003). Together with our data, these findings suggest potential biological relevance of ALDH3A1 in ESCC development which warrants further investigations.

To exploit the epigenomic dysregulation in ESCC for potential therapeutic intervention, we investigated both BET and HDAC inhibitors. BET inhibition was tested because *BRD4* was highly expressed in ESCC and was identified as a direct target of ESCC CRC members. We also discovered that, a HDAC inhibitor (Romidepsin) exhibited prominent capabilities of degrading TP63 protein, and consequently destructing the entire ESCC CRC (Figure 8C). Not surprisingly, either ARV-771 (BET degrader) or Romidepsin elicited pronounced anti-neoplastic effect in ESCC cells, with ensuing reduction of H3K27ac and H3K4me3 signals across the genome. ARV-771 and Romidepsin co-targeted thousands of shared regulatory regions. Moreover, these two compounds exhibited differential and complementary effects on the ESCC epigenome, with ARV-771 preferentially impairing promoter functions while Romidepsin favoring the inhibition of distal enhancers. Supporting our data, HDAC inhibitors attenuated acetylation at distal regulatory elements and decreased expression of associated genes in colorectal and pancreatic cancer cells (Mishra et al., 2017; Sanchez et al., 2018). In the present study, the function of Romidepsin on distal enhancers is likely to be attributed to the potency of degradation of TP63 as evidenced by the shared changes in enhancer loci (Figure 8G). Based on their differential and complementary effects on the ESCC epigenome, we tested and confirmed a synergistic anti-ESCC effect when ARV-771 and Romidepsin were combined both *in vitro* and *in vivo*. Likewise, the enhanced anti-proliferative activity of a BET protein with an HDAC was demonstrated as a potential therapeutic strategy for both pancreatic cancer (Mazur et al., 2015) and *Myc*-induced murine lymphoma (Bhadury et al., 2014). Together, this combinatorial strategy deserves further pre-clinical and clinical testing for ESCC treatment.

## STAR**★**METHODS

Detailed methods are provided in the online version of this paper and include the following:

- KEY RESOURCES TABLE
- CONTACT FOR REAGENT AND RESOURCE SHARING
- EXPERIMENTAL MODEL AND SUBJECT DETAILS

- Animal Studies
- Human Cell Lines
- METHOD DETAILS

- Chromatin Immunoprecipitation
- ATAC-seq
- Data Analysis for ChIP-seq and ATAC-seq
- Annotation of Super-Enhancers (SEs) and Establishment of CRC Program
- Circular Chromosome Conformation Capture sequencing (4C-seq) Assay
- Chromosome Conformation Capture (3C) Assay
- CRISPR/Cas9-mediated Genomic Deletion of Regulatory DNA Regions
- RNA-seq Analysis
- Western Blotting and Immunoprecipitation
- Cell Viability Assay and Colony Formation Assay
- Transfection of siRNAs and Plasmids
- Lentiviral Production and Generation of Stable Cell Lines
- QUANTIFICATION AND STASTICAL ANALYSIS
- DATA AND SOFTWARE AVAILABILITY

## SUPPLEMENTAL INFORMATION

Supplemental Information includes seven figures and nine tables.

## ACKNOWLEDGMENTS

We thank Dr. Takashi Yamamoto (Department of Mathematical and Life Sciences, Graduate School of Science, Hiroshima University, Japan) for sharing the all-in-one CRISPR/Cas9 vectors. This work is funded by the Singapore Ministry of Health’s National Medical Research Council (NMRC) under its Singapore Translational Research (STaR) Investigator Award to H. Phillip Koeffler (NMRC/STaR/0021/2014), Singapore Ministry of Education Academic Research Fund Tier 2 (MOE2013-T2-2-150), the NMRC Centre Grant Programme awarded to National University Cancer Institute of Singapore (NMRC/CG/012/2013 and CGAug16M005) and the National Research Foundation Singapore and the Singapore Ministry of Education under its Research Centres of Excellence initiatives.

## AUTHOR CONTRIBUTIONS

Y.Y.J., Y.J., and D.C.L. conceptualized and devised the study. Y.Y.J., and Y.J. designed experiments and analysis, and H.K., and J.W.D helped perform the experiments. Y.Z., and L.A. performed 4C-seq and 3C analysis. Y.Y.J., C.Q.L, P.D., A.M., and Y.Y.L. performed bioinformatics and statistical analysis. R.Y.T.L. helped establish xenograft mice. L.H., J.J.X., M.H., L.X., M.J., D.K., and X.Y.G. contributed resources or critical feedback on project. Y.Y.J., Y.J., M.J.F., D.C.L., and H.P.K. analyzed the data. M.J.F., D.C.L., and H.P.K. supervised the research. Y.J., and Y.Y.J. wrote original draft, together with D.C.L., and H.P.K. reviewed and edited the manuscript.

## Supplement Figures

**Figure S1.**
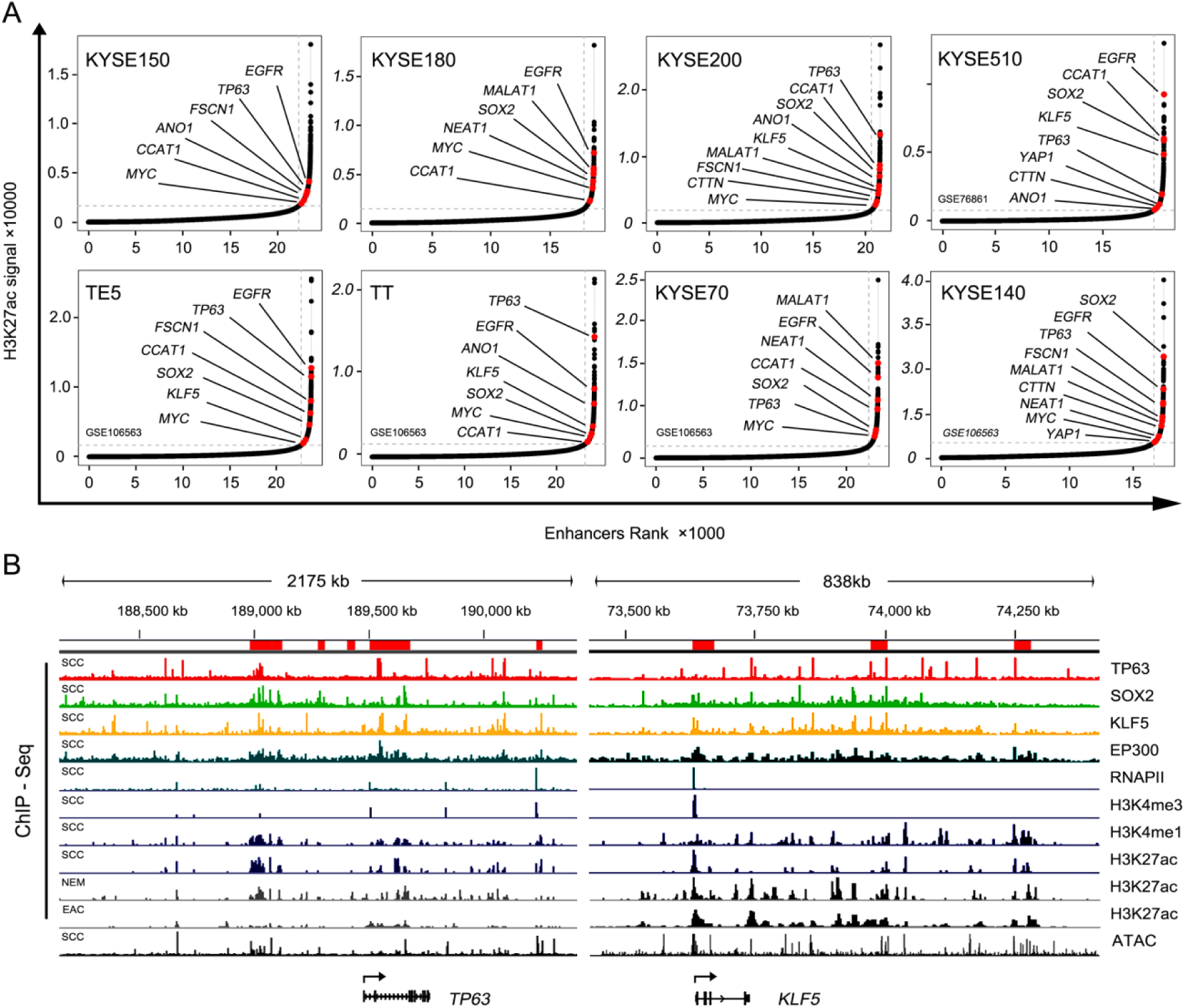
Super-enhancer (SE) Profiles of ESCC cells. Related to Figure 1. (A) Hockey stick plots based on input-normalized H3K27ac ChIP-seq signal identifying super-enhancers (SEs) in 8 ESCC cell lines. X axis ranked by enhancer regions; Y axis plotted H3K27ac enrichment. (B) Integrative genomics viewer (IGV) showing normalized ChIP-seq signals for indicated antibodies and ATAC-seq tracks at the *TP63* and *KLF5* loci in squamous cell carcinoma (SCC), esophageal adenocarcinoma (EAC) and nonmalignant esophageal mucosa (NEM) samples.

**Figure S2.**
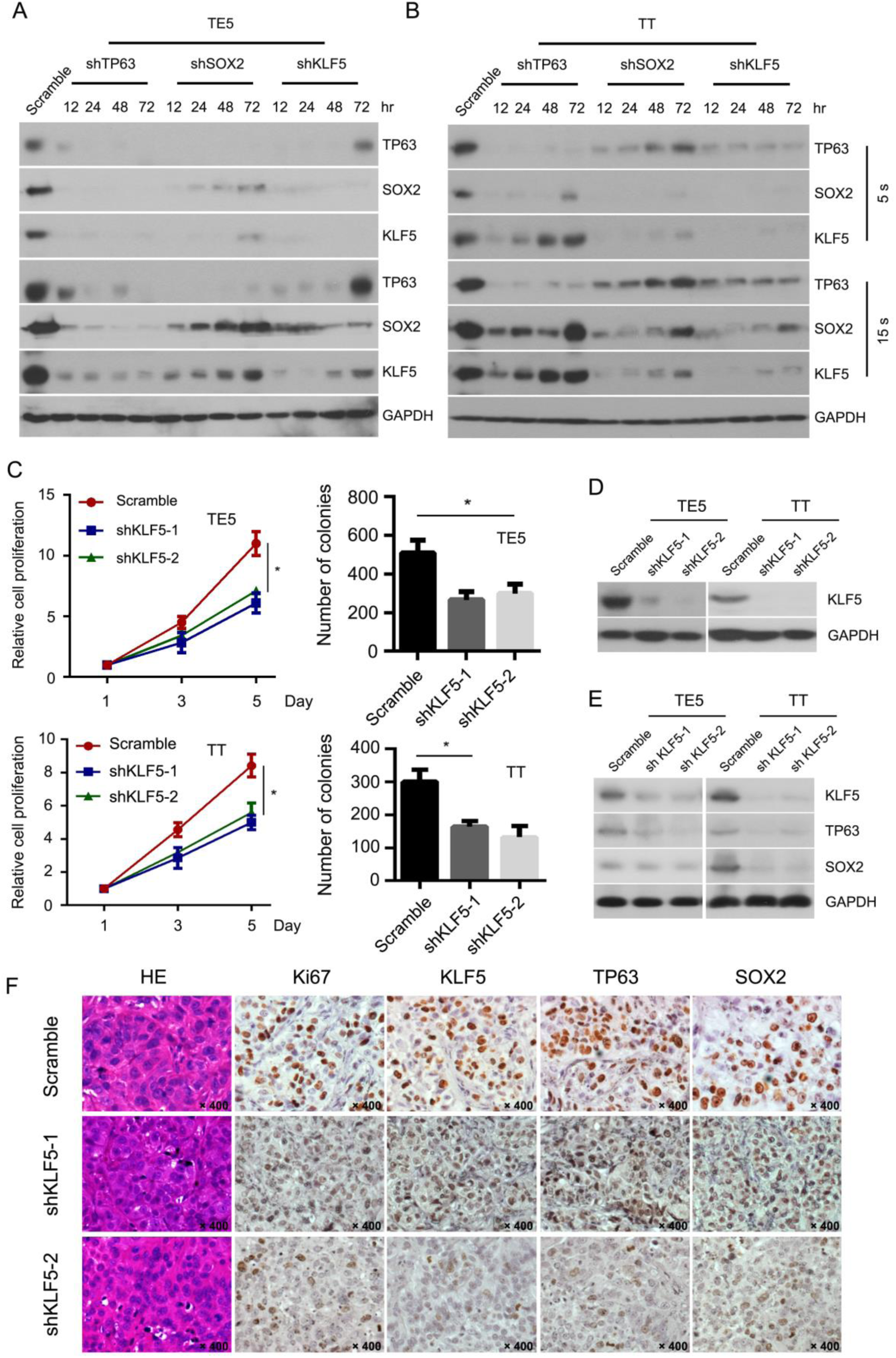
Interplay of TP63/SOX2/KLF5 and Oncogenic Property of *KLF5* in ESCC. Related to Figure 2. (A and B) Western blotting arrays presenting expression of TP63, SOX2, KLF5 by separately downregulation of each TF at indicated time-course with 5 seconds (s) and 15 seconds (s) exposure time in TE5 (A) and TT (B) cells. (C) Cell viability measured by both MTT assay (Left) and colony formation assay (Right) upon silencing of KLF5 with two independent shRNAs in both TE5 and TT cells. Error bars, mean ± SD. * P < 0.05. (D) Western blotting showing the efficient downregulation of KLF5 with two independent shRNA targets. (E) Western blotting array measuring expression of KLF5 and the two other TFs of CRC in ESCC xenografts either with or without knockdown of KLF5. (F) H&E and IHC staining of murine tumors for cell proliferation marker (Ki67), and CRC TFs in either scramble or KLF5-knockdown cells-derived xenografts. Original magnification: × 400.

**Figure S3.**
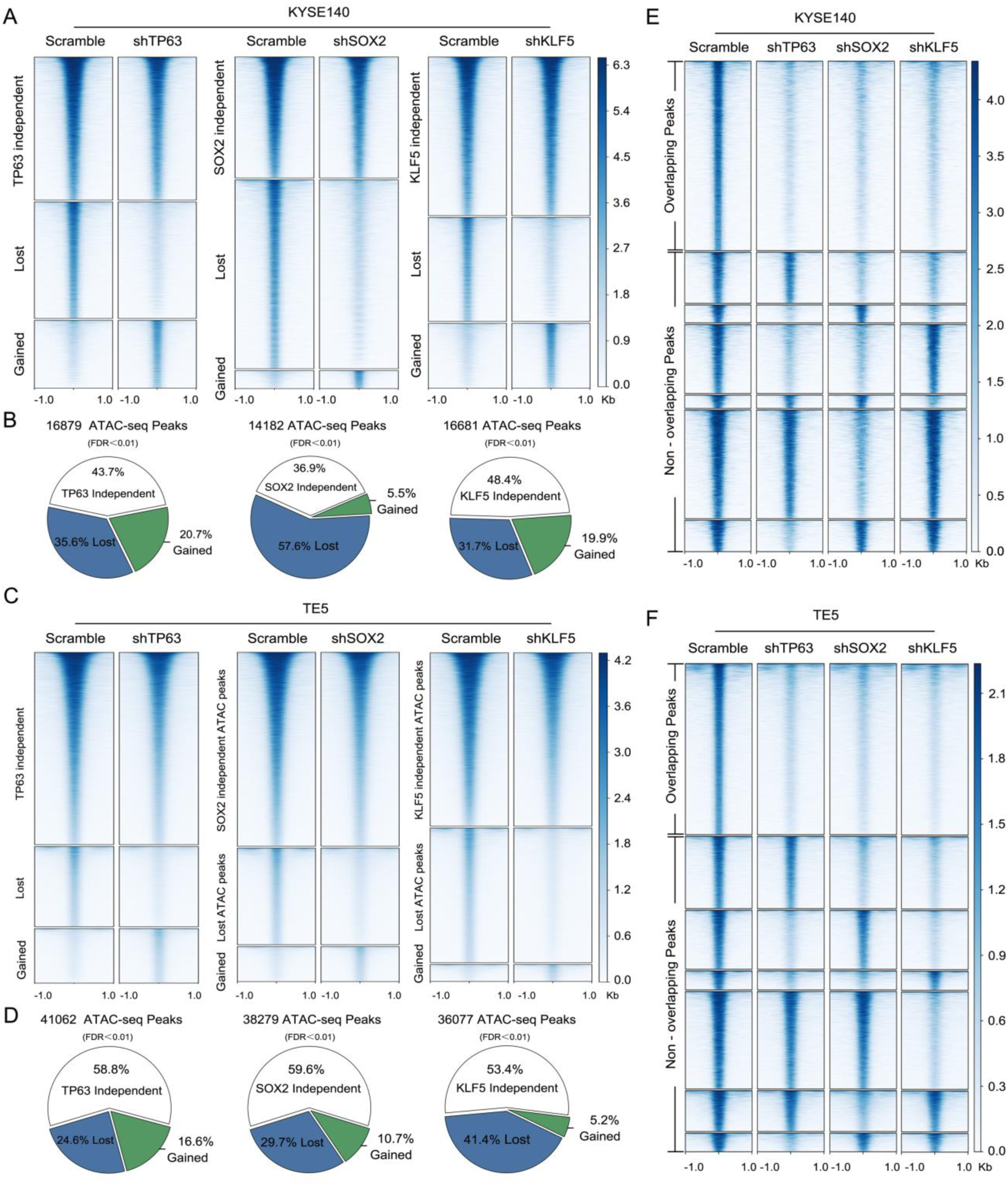
Altered Chromatin Accessibility Caused by Silencing of TP63, SOX2 or KLF5. Related to Figure 3. (A-D) Heatmaps (A and C) and pie charts (B and D) showing proportion of altered chromatin accessibility with downregulation of either TP63, SOX2 or KLF5 in KYSE140 (A and B) and TE5 cells (C and D). (E and F) Heatmaps of lost ATAC-seq signals upon downregulation of each of TF in KYSE140 (E) and TE5 (F) cells in comparison to scramble control. These regions are further stratified by the degree of overlapping among each experiment.

**Figure S4.**
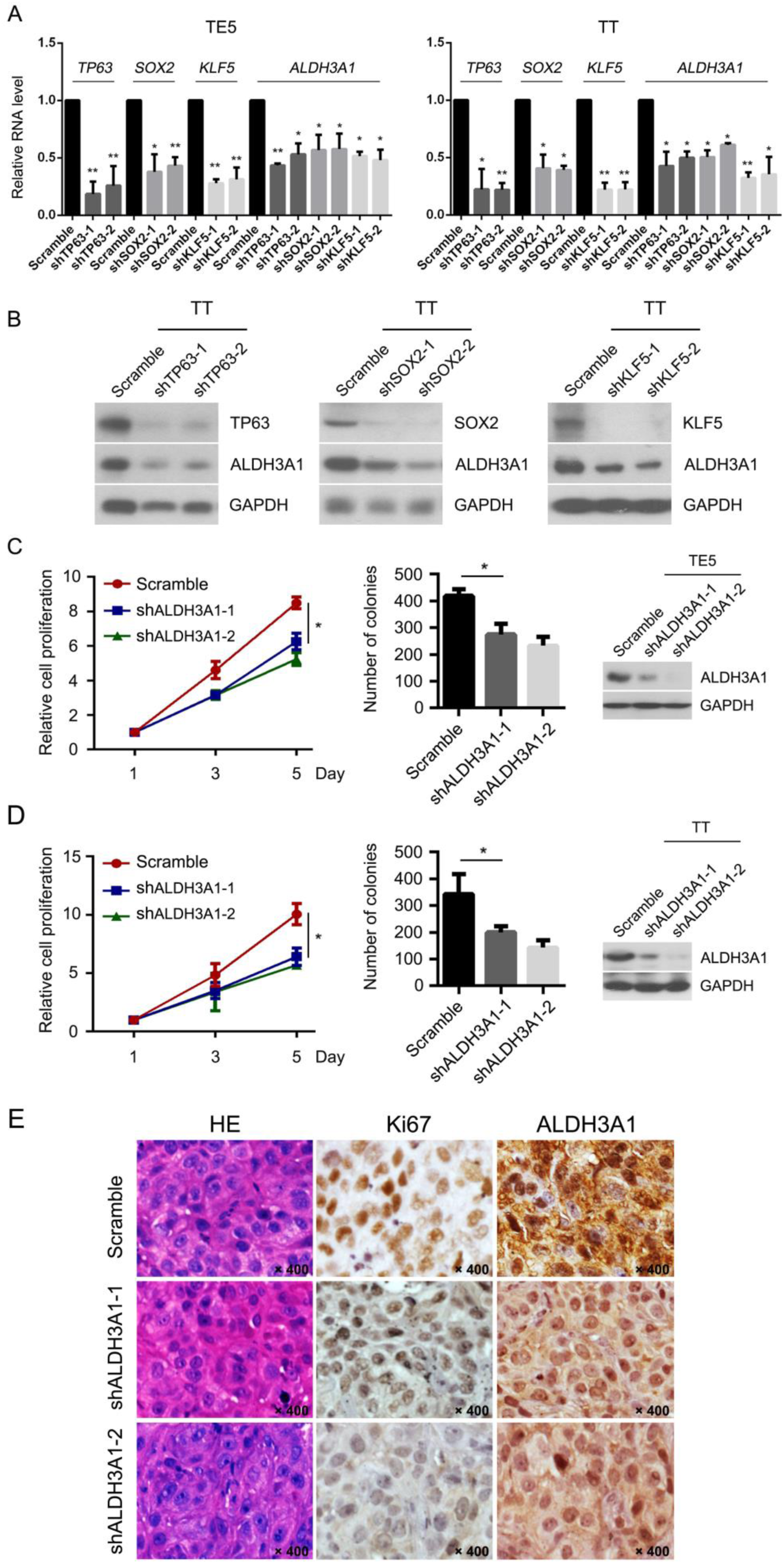
*ALDH3A1* as a SE-associated Oncogene is Directly Regulated by CRC TFs in ESCC. Related Figure 4. (A) Relative mRNA expression of *ALDH3A1* upon silencing of either TP63, SOX2 or KLF5 in TE5 and TT cell lines. Data represent means ± SD. * P < 0.05, ** P < 0.01. (B) Western blotting showing protein levels of ALDH3A1 in TT cells upon silencing of each of CRC TF. (C and D) MTT assay (Left) and colony formation assay (Middle) upon silencing of ALDH3A1 in TE5 (C) and TT (D) cell lines. Right, Western blotting verifies efficient knockdown of ALDH3A1 with two independent shRNAs. Data represent means ± SD. * P < 0.05. (E) H&E and IHC staining of xenograft tumor sections, IHC measures the cell proliferation marker (Ki67) as well as ALDH3A1. Original magnification: × 400.

**Figure S5.**
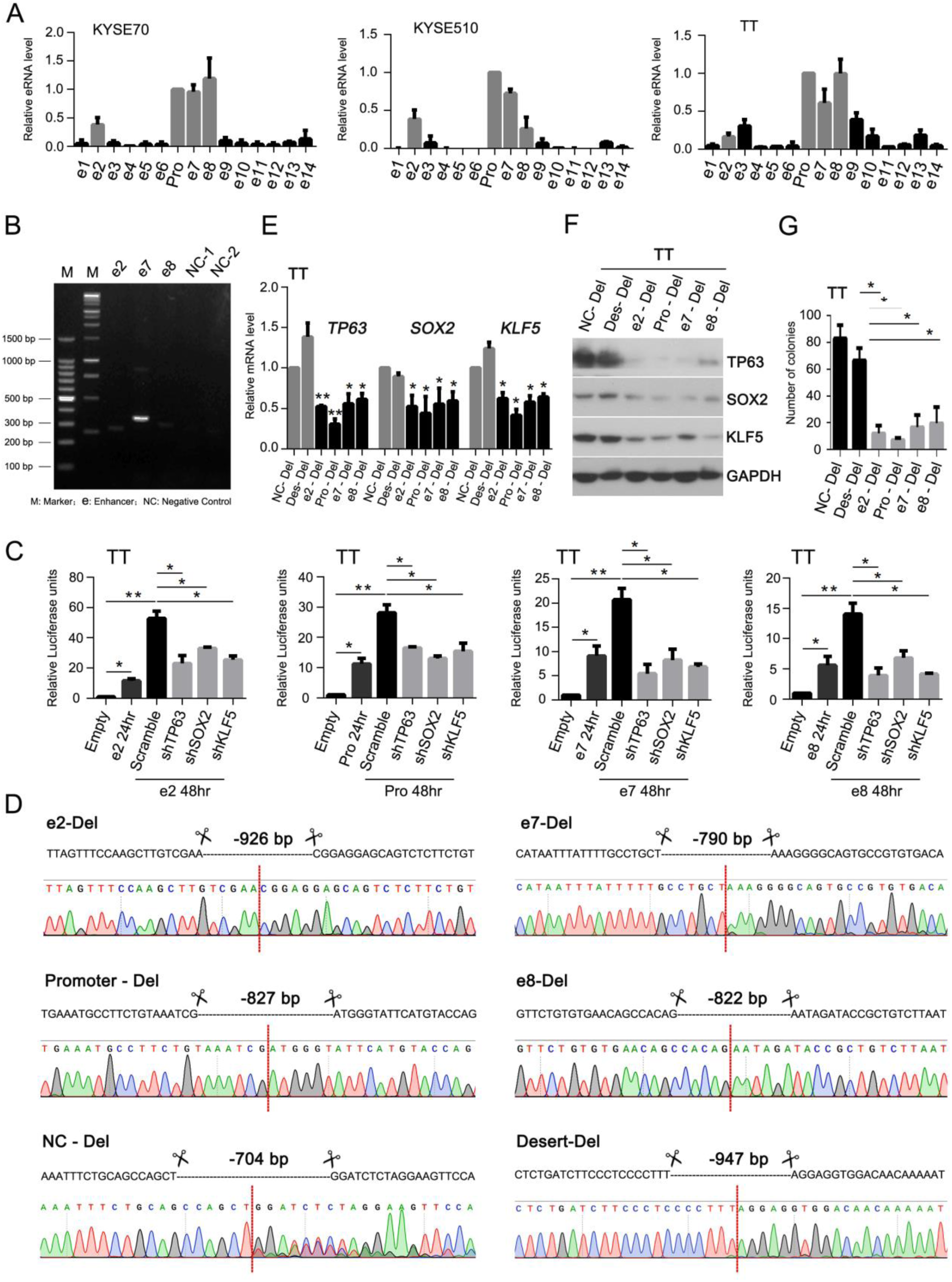
Interdependence between *TP63* Essential Enhancers and CRC TFs. Related to Figure 6. (A) Relative eRNA expression of each of enhancer component in KYSE510, KYSE70 and TT cell lines. Data are normalized to RNA expression of the promoter. Pro: Promoter. (B) Agarose gel electrophoresis following 3C analysis showing interaction between *TP63* promoter and enhancers: e2, e7 and e8. NC1 and NC2 are negative controls (elements showing no interaction with *TP63* promoter according to 4C-Seq analysis). NC (Negative Control). (C) Box plots showing relative luciferase activities in TT cells either with or without silencing of TP63, SOX2 or KLF5. (D) Representative Sanger sequencing for each region with CRISPR/Cas9 and paired sgRNAs-mediated deletion. Red dotted line indicates the rejoining ends. Del: deletion. (E-G) Statistical analysis of mRNA expression (E), protein levels of CRC TFs (F) and colony number (G) in unedited control and edited TT cells. NC and Des (Desert) serve as negative controls. Data of (C), (E), and (G) represent the mean ± SD. * P < 0.05, ** P < 0.01.

**Figure S6.**
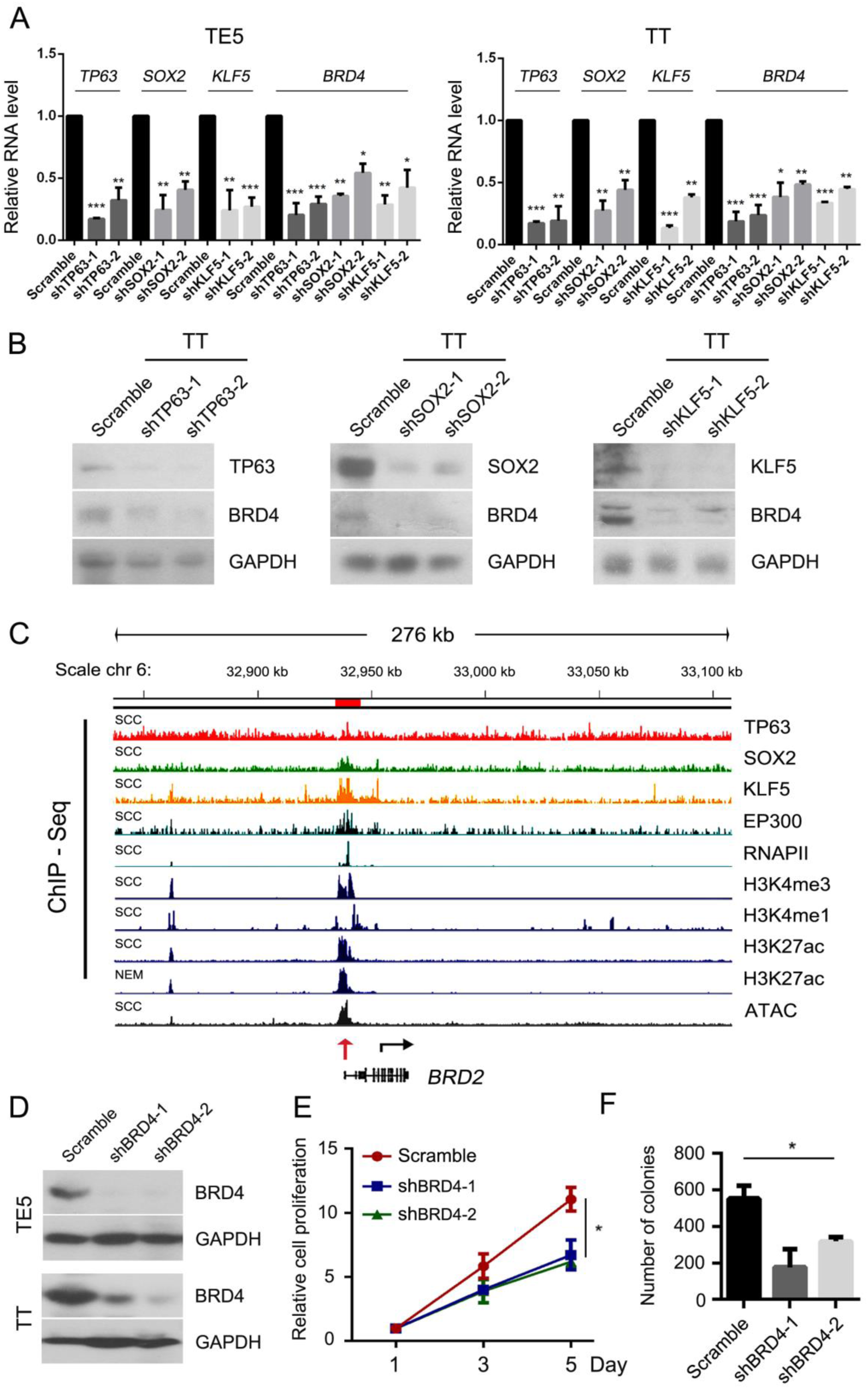
Necessity of BRD4 for ESCC Growth. Related to Figure 7. (A and B) qRT-PCR and Western blotting showing relative levels of BRD4 mRNAs (A) and proteins (B) upon silencing of either TP63, SOX2 or KLF5 in TE5 and TT cells. (C) IGV tracks of ChIP-seq for indicated antibodies and ATAC-seq around *BRD2* gene locus. (D) Western blotting showing efficient knockdown of BRD4 at protein level with two independent shRNAs in TE5 and TT cells. (E and F) MTT (E) and colony formation (F) assays upon silencing of BRD4 in TT cells.

**Figure S7.**
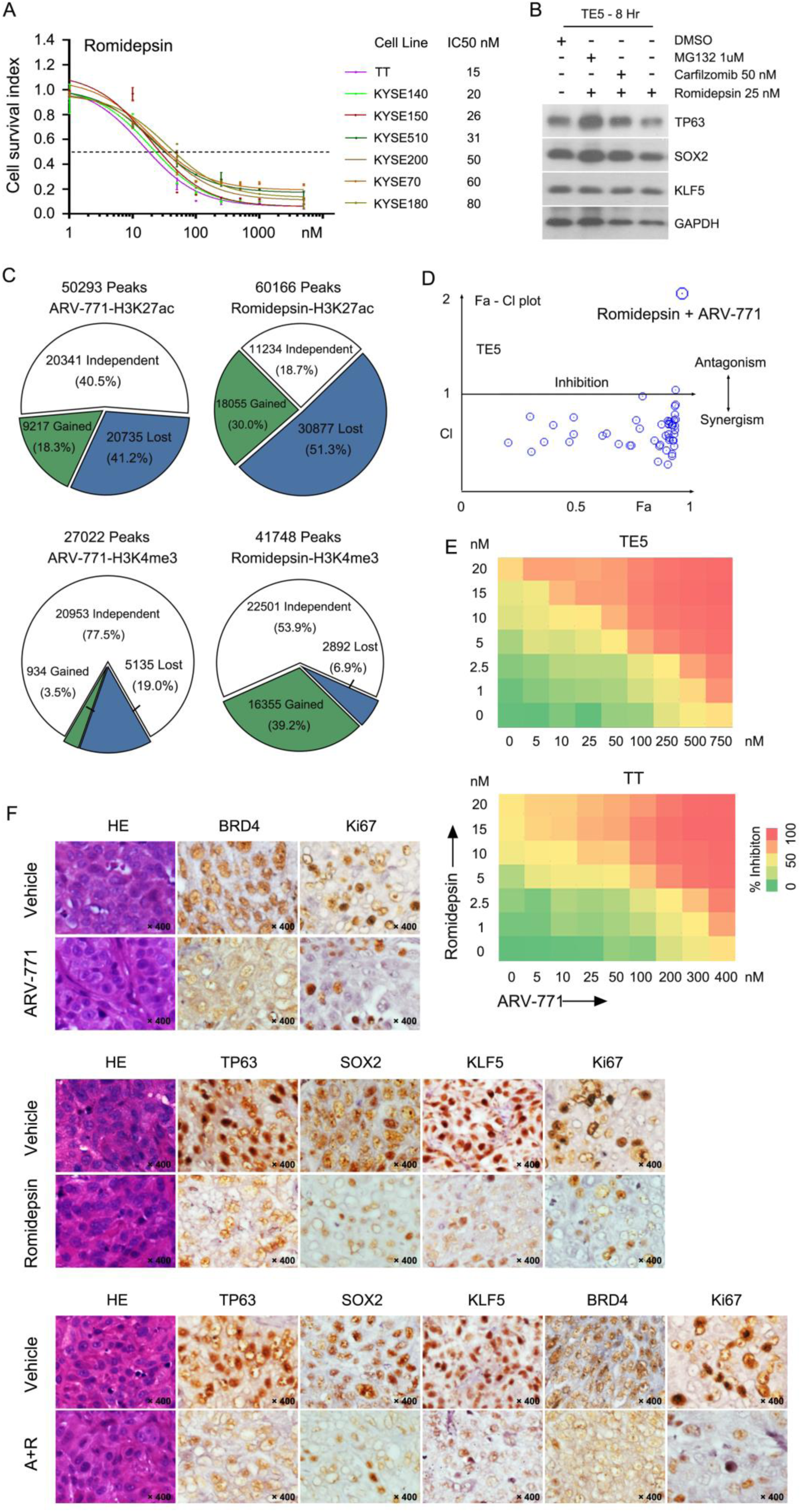
HDAC and BET Inhibition Synergistically Represses ESCC Cell Growth. Related to Figures 8. (A) Cellular viability curves of ESCC cell lines after exposure to different concentration of Romidepsin (3 days), measured by MTT. IC50 is shown on the right. (B) Western blotting array showing protein levels of TP63, SOX2 and KLF5 in TE5 cells upon treatment of indicated drugs. (C) Pie chart showing the changed proportion of H3K4me3 and H3K27ac peaks upon the treatment with either ARV-771 or Romidepsin using TT cells. (D) Fa-CI plots showing synergistic inhibition of ARV-771 and Romidepsin using TE5 cells, calculated according to results from (E). CI < 1, = 1, and > 1 indicate drug synergism, additive effect, and antagonism, respectively. (E) Heatmap matrix showing percent inhibition of TT cells with combinatorial treatment with either ARV-771 or Romidepsin. Cell viability was measured by MTT on 3 day. Yellow colors represent 50% inhibition. (F) H&E and IHC staining of tumor sections. IHC measured Ki67, TP63, SOX2 and KLF5 in xenograft tumors after administration of either vehicle, ARV-771, Romidepsin or both. Original magnification: × 400.

